# Quality matters: stoichiometry of resources modulates spatial feedbacks in aquatic-terrestrial meta-ecosystems

**DOI:** 10.1101/2023.03.15.532732

**Authors:** Benôıt Pichon, Elisa Thébault, Gérard Lacroix, Isabelle Gounand

## Abstract

Species dispersal and resource spatial flows greatly affect the dynamics of connected ecosystems. So far, research on meta-ecosystems has mainly focused on the quantitative effect of subsidy flows. Yet, resource exchanges at heterotrophic-autotrophic (*e.g.,* aquatic-terrestrial) ecotones display a stoichiometric asymmetry that likely matters for functioning. Here, we joined ecological stoichiometry and the meta-ecosystem framework to understand how subsidy stoichiometry mediates the response of the meta-ecosystem to subsidy flows. Our model results demonstrate that resource flows between ecosystems can induce a positive spatial feedback loop, leading to higher production at the meta-ecosystem scale by relaxing local ecosystem limitations (“spatial complementarity”). Furthermore, we show that spatial flows can also have an unexpected negative impact on production when accentuating the stoichiometric mismatch between local resources and basal species needs. This study paves the way for studies on the interdependancy of ecosystems at the landscape extent.

**Data:** The code and the data, as well as a small tutorial to run the model are available on Github via Zenodo: https://doi.org/10.5281/zenodo.7733880

## Introduction

Flows of organisms, resources, and energy connect communities and ecosystems at the landscape scale (Polis <u>et al</u>., 1997). These spatial connections are key to consider for understanding ecosystem functioning and its response to global changes. As habitats are increasingly fragmented, species indeed often disperse to track more favorable abiotic conditions (Haddad <u>et al</u>., 2015; Thompson & Fronhofer, 2019). In addition, ongoing changes in resource flows between ecosystems greatly affect ecosystem functioning, as exemplified by the consequences of increasing terrestrial organic matter inputs on aquatic ecosystems (Solomon <u>et al</u>., 2015). In this context, the meta-ecosystem framework (*sensu* Loreau <u>et al</u>., 2003) unveils the strong links between local processes and landscape dynamics (Polis <u>et al</u>., 2004; Massol <u>et al</u>., 2011). Meta-ecosystem theory has shown that spatial flows at the landscape scale can affect source-sink dynamics (Gravel <u>et al</u>., 2010), the local coexistence of communities (Marleau & Guichard, 2019; Peller <u>et al</u>., 2021), and both the stability and functioning of ecosystems (Marleau <u>et al</u>., 2014; Gounand <u>et al</u>., 2014). However, this theory has so far been restricted to flows between identical ecosystems mediated by species dispersal. It has focused on flow magnitude, ignoring the importance of flow quality (but see Marleau <u>et al</u>., 2015). While adapted to describe spatial networks of connected forest patches or lakes, these models do not capture the diversity of resource flows crossing ecotones, such as terrestrial-aquatic ecotones (Massol <u>et al</u>., 2017; Gounand <u>et al</u>., 2018a).

Indeed, resource flows (*i.e.*, detritus or nutrients) of varying quantity and quality cross ecotones and connect dissimilar ecosystems (*e.g.*, forest and streams), with strong impacts on recipient ecosystems (Bartels <u>et al</u>., 2012). For instance, plant litter fuels stream communities with carbon-rich subsidies (Wallace <u>et al</u>., 1997; Nakano & Murakami, 2001; Bartels <u>et al</u>., 2012) while fish caught from streams by terrestrial predators feed riparian communities with nutrient-rich carcasses, providing as much as 25% of the nitrogen budget in riparian forests (Baxter <u>et al</u>., 2005; Helfield & Naiman, 2006). A recent data synthesis showed that carbon resources commonly flow from net autotroph ecosystems (*i.e.*, where primary production exceeds ecosystem respiration), to net heterotroph ones, where primary production is limited by different factors (*e.g.*, shade, aridity, or water depth in benthic systems; Gounand <u>et al</u>., 2018b). In many lakes and small shaded streams, for instance, more carbon is respired than fixed, making their functioning net-heterotroph and dependent on terrestrial carbon flows (Bartels <u>et al</u>., 2012; Gounand <u>et al</u>., 2018b). By contrast, terrestrial or pelagic net autotrophic systems receive smaller carbon flows, which seem negligible compared to their primary production (Gounand <u>et al</u>., 2018b). This asymmetry in the quantity of resources transferred between autotroph and heterotroph ecosystems can, however, be balanced by a stoichiometric asymmetry. Here we complemented the carbon view of spatial coupling at terrestrial-freshwater ecotones by data on nitrogen spatial flows (Appendix S2): we show that the asymmetry of resource flows is reversed in terms of quality of resources (*i.e.*, N:C ratio), leading to nitrogen spatial flows of the same order of magnitude to aquatic and terrestrial ecosystems (Fig. 1, Box 1). Indeed, organisms produce detritus of different qualities, depending on their stoichiometric composition: resource flows from terrestrial ecosystems are mostly composed of primary producers with low N:C ratios, whereas the resource flows from freshwater ecosystems originate from organisms at higher trophic levels with higher N:C ratios (Fig. 1, Elser <u>et al</u>., 2000; Sitters <u>et al</u>., 2015). Therefore, subsidy stoichiometry likely plays a strong role in the functioning of aquatic-terrestrial meta-ecosystems, as already suggested at the ecosystem level.

**Figure 1:**
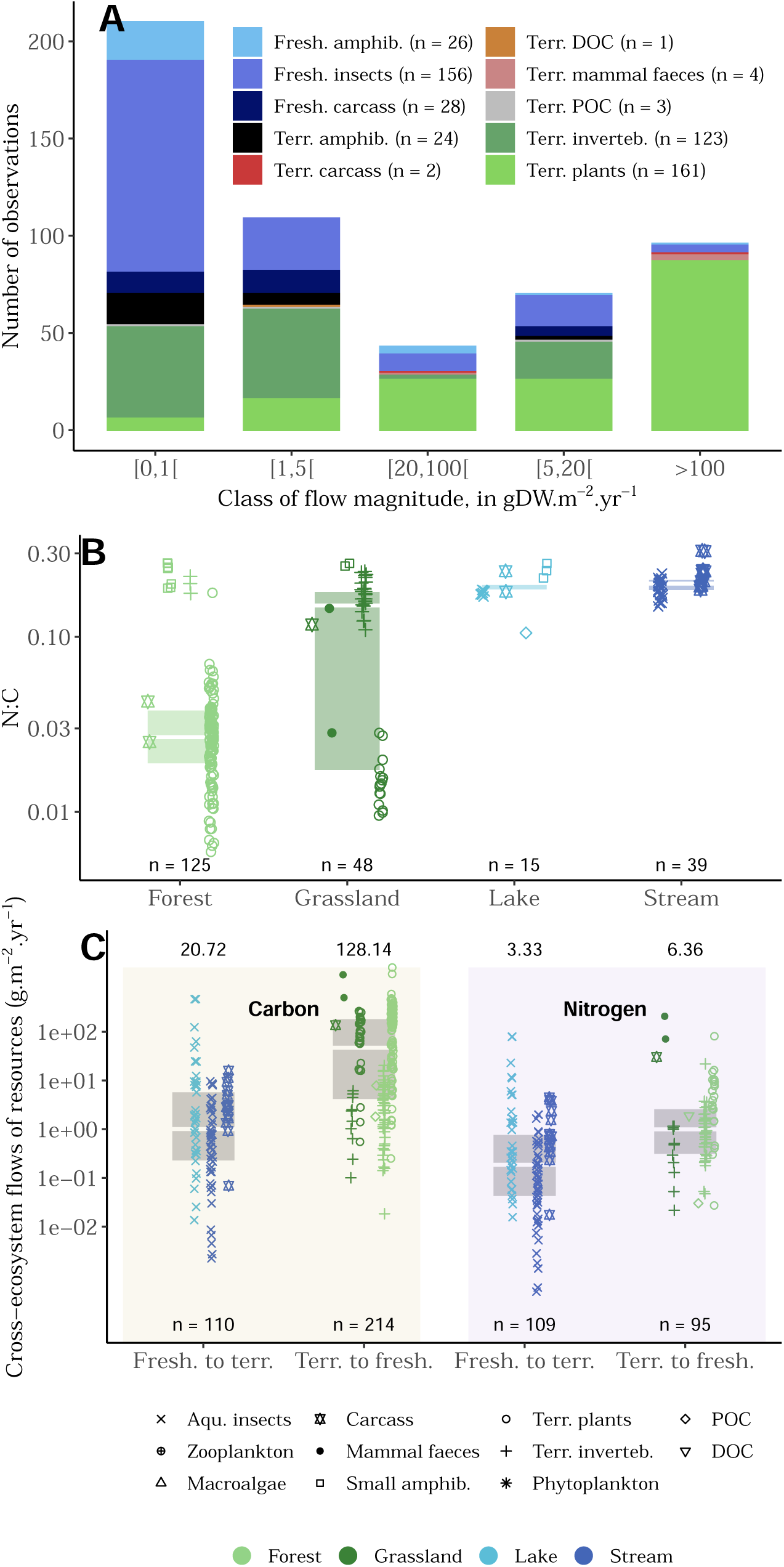
Empirical data on stoichiometry and magnitude differences between resources exchanged at freshwater-terrestrial ecotone. (A): Distribution of the number of observations of cross-ecosystem flows at terrestrial-freshwater ecotone according to their magnitude (in *gDW*.*m*^−2^.*yr*^1^, see Appendix S2 for data extraction). The surface (in *m^−^*^2^) corresponds to the area of the ecosystem receiving the resource flow. (B): Nitrogen to carbon ratio (molar) of materials exported by terrestrial (forests and grasslands in green) and freshwater (streams and lakes in blue) ecosystems. The shape of the points indicates the type of materials being exported by ecosystems. Fresh. = Freshwater, Terr. = Terrestrial, amphib. = amphibians and inverteb. = invertebrates. Terrestrial or aquatic labels indicate the donor ecosystem. (C): Cross-ecosystem flows of resources (carbon (left) and nitrogen (right)) crossing freshwater-terrestrial ecotone. The units of flows are *gC*.*m*^−2^.*yr*^1^ and *gN*.*m*^−2^.*yr*^1^ for carbon and nitrogen respectively. Fresh. = Freshwater, Terr. = Terrestrial. Boxplots present the median surrounded by the first and third quartiles, and whiskers are not shown. Mean flows are indicated on the top of each boxplot. Note that, panel (B) shows the stoichiometry of material **exported** by each type of ecosystem, while panel (C) represents the quantity of C and N fluxes fueling each type of ecosystem. The number of observations, *n*, is indicated in each panel.

Differences in stoichiometry and limitations between organisms in local communities are known to strongly impact local ecosystem processes ranging from nutrient recycling by consumers to species coexistence (Daufresne & Loreau, 2001; Cherif & Loreau, 2013; Daufresne, 2021). The stoichiometry of spatial flows exported is thus expected to affect the functioning of connected ecosystems, depending on the limitation of their basal species. Primary producers are mainly limited by nutrients (*i.e.,* phosphorus or nitrogen) in autotrophic systems, while in heterotrophic systems where primary production is low, species at the basis of the food web are decomposers that can also be limited by carbon (Elser <u>et al</u>., 2007; Daufresne <u>et al</u>., 2008; Harpole <u>et al</u>., 2011). Changing the limitation of organisms within an ecosystem can drastically change the interactions and the feedbacks between trophic levels (Zou <u>et al</u>., 2016; Buchkowski <u>et al</u>., 2019). Together, this suggests that the impact of subsidy flows on ecosystem and meta-ecosystem scale production may depend on both *(i)* the stoichiometry of subsidies and *(ii)* the local stoichiometric constraints on communities.

Recent experimental and theoretical works suggest a functional complementarity of ecosystems at the landscape scale (Gounand <u>et al</u>., 2017; Harvey <u>et al</u>., 2021). For instance in watersheds, terrestrial ecosystems generally drive autotrophic production at the landscape extent while some aquatic ecosystems, such as rivers with high riparian cover or lakes, could display net heterotrophic functioning and perform intensive decomposition processes (Gounand <u>et al</u>., 2018b). In addition, aquatic ecosystems tend to have higher trophic efficiency (*i.e.,* ratio of consumer to prey production; Shurin <u>et al</u>., 2006), and may boost landscape-scale secondary production using terrestrial carbon resources. Thus, resource flows may optimize different functions in the meta-ecosystem (Harvey <u>et al</u>., 2021). Studies on aquatic-terrestrial meta-ecosystems have so far considered differences in trophic efficiency and primary production (Evans-White & Halvorson, 2017). We hypothesize that similar mechanisms can be expected with stoichiometry. If spatial flows fuel local ecosystems with resources that are limiting producer growth (*e.g.,* carbon-rich subsidies from terrestrial ecosystems relaxing carbon limitation of decomposers in streams), we expect higher production and positive feedbacks to emerge at the meta-ecosystem scale. However, to our knowledge, no theoretical and empirical studies investigated such hypotheses.

Here we integrate ecological stoichiometry and the meta-ecosystem framework to understand *(i)* how stoichiometry mediates the response of meta-ecosystems to subsidy flows and *(ii)* whether subsidy flows increase meta-ecosystem production by inducing positive spatial feedback through the relaxation of local limitation in carbon or nutrients. To test our hypotheses, we develop a stoichiometrically explicit meta-ecosystem model, which connects the dynamics of a heterotrophic and an autotrophic ecosystem by carbon and nitrogen fluxes (*e.g.,* aquatic-terrestrial ecotones). To root our narrative in natural systems, and since we focused our empirical data search on terrestrial-freshwater ecotones, we hereafter assign aquatic and terrestrial labels to our modelled ecosystems, but our framework is generic to any autotrophic-heterotrophic meta-ecosystem and could, for example, be applied to benthic-pelagic dynamics. With this model, we explore how flows of carbon and nitrogen subsidies interact with local community dynamics to determine meta-ecosystem functioning under different scenarios of resource limitations. Our results reveal non-linear positive feedbacks between ecosystems under carbon limitation of the aquatic ecosystem, but negative feedbacks under nitrogen limitation. Spatial flows of resources increase or decrease overall production depending on whether they accentuate the stoichiometric complementarity or the mismatch, respectively, between the needs of local communities and the composition of available resources.

## Material and methods

### Model description

#### Meta-ecosystem model

We built a meta-ecosystem model with explicit stoichiometry that couples carbon and nitrogen dynamics between two ecosystems with contrasting functioning (Fig. 2): one autotroph, hereafter labelled terrestrial (indexed by *T*, *e.g.,* forest or grassland) and one heterotroph, hereafter labelled aquatic (indexed by *A*, *e.g.,* stream with a high riparian cover or oligotrophic lake). We focus on the autotrophic-heterotrophic extrema of a gradient in ecosystem functioning, by considering the basal species of the terrestrial ecosystem to be autotrophic plants (*P*), while the aquatic ecosystem harbors no primary producers, and instead has heterotrophic decomposers (*B*) as basal species. We assume that each ecosystem consists of two trophic levels to ease the interpretation of our results. While spatial flows exported by aquatic ecosystems mainly originate from higher trophic levels (Fig. 1), this assumption does not affect our results (see Appendix S4 for results with an additional trophic level). Therefore, grazers (*G*) and consumers of decomposers (*C*) consume basal species in terrestrial and aquatic ecosystems, respectively. Each ecosystem has a detritus (*D*) and an inorganic nitrogen pool (*N*). We followed the carbon for the biotic and detritus pools (*e.g., B_CA_* for aquatic decomposers), and we derived the nitrogen content of organisms (*e.g.*, *P_N_* for plants) by assuming a fixed stoichiometry (homeostasis) corresponding to their molar nitrogen to carbon ratio, N:C (*α_X_*, where *X ∈ {G*, *P*, *C*, *B}* ; Fig. 2A). This homeostasis involves different mechanisms across trophic levels: contrary to plants that maintain a constant N:C ratio through the control of their uptake (Tilman, 1982), grazers and consumers of decomposers hold their homeostasis through differential assimilation (Grover & Holt, 1998; Sterner & Elser, 2002).

**Figure 2:**
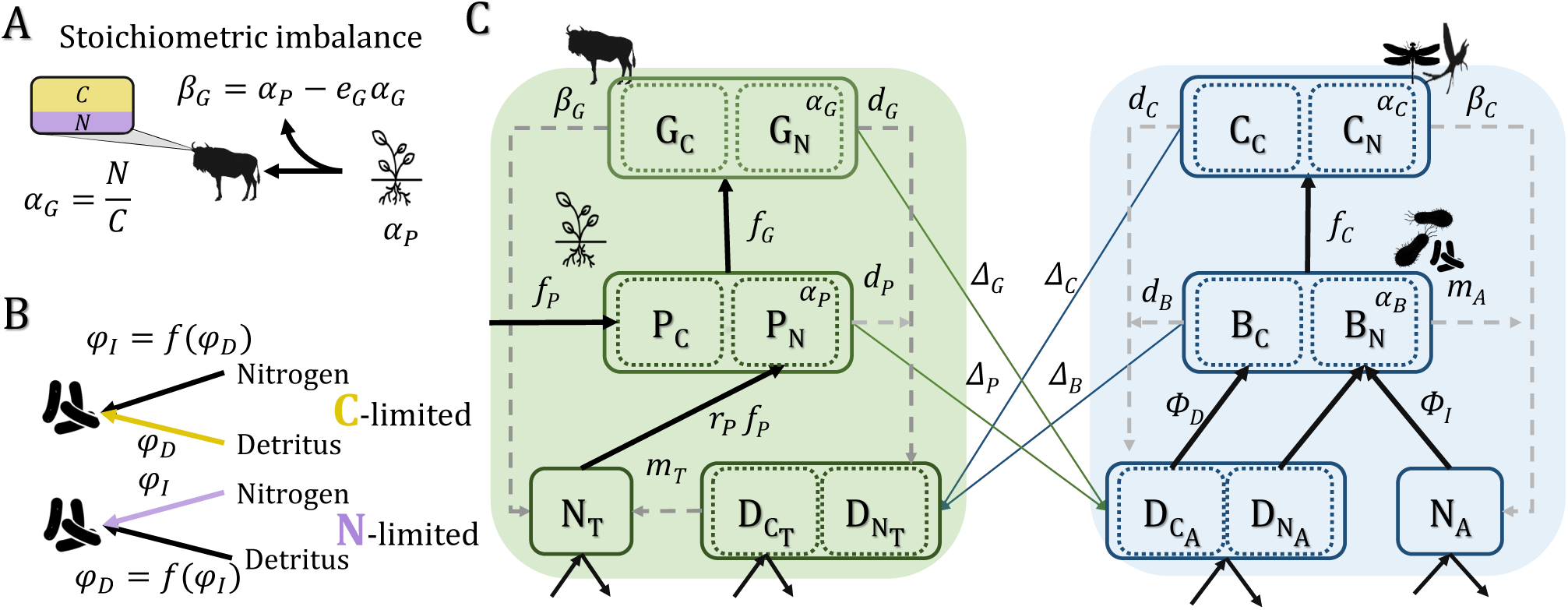
Stoichiometric meta-ecosystem model - hierarchy of processes and scales. Basal species and consumers are held at a fixed stoichiometry (homeostasis). (A): Stoichiometric imbalance (*β_G_*) results from the differences between producer stoichiometry and consumer needs and lead to nitrogen excretion. A similar mechanism occurs for the interaction between decomposers and their consumers. (B): Decomposers from the freshwater heterotrophic ecosystem have two possible limitations: nitrogen or carbon. Their limitation sets which flow (decomposition of detritus (*ϕ_D_*) or immobilization of nitrogen (*ϕ_I_*)) constrains the other so that decomposers maintain a constant stoichiometry. (C): Two ecosystems, one heterotrophic (freshwater) with decomposers as basal species and the other autotrophic (terrestrial) with plants as primary producers, are linked at the landscape level by subsidy flows (dead organic matter and nitrogen flows due to stoichiometric imbalance). Symbols are used in the model equations: mineralization rates in ecosystems (*m_T_*, *m_A_*), immobilization flow (*ϕ_I_*), decomposition flow (*ϕ_D_*), plant net photosynthesis (*f_P_*) and nitrogen uptake (*α_P_ f_P_*), attack rates of consumers of decomposers and grazers (*f_C_*, *f_G_* respectively), decay rates of organisms (*d_G_*, *d_P_*, *d_B_*, *d_C_*), the stoichiometric imbalance between producers and consumers (*β_G_*, *β_C_*), the fraction of subsidies recycled regionally (Δ*_P_*, Δ*_G_*, Δ*_C_*, Δ*_B_*) and nitrogen to carbon ratio of organisms (*α_G_*, *α_C_*, *α_P_*, *α_B_*) Black thick arrows describe trophic interactions while grey dashed arrows correspond to recycling paths.

#### Basal species limitation

Plants are exclusively limited by nitrogen for growth and maintain their stoichiometric homeostasis by adjusting their carbon intake, such that net photosynthesis equals *f_P_* (Eq 1). Decomposers can either be limited by carbon or nitrogen (Fig. 2B, or by both in Appendix S5). Decomposers feed primarily on detritus. Yet, as the detritus N:C ratio does not always match the decomposer stoichiometry due to different stoichiometry of detritus produced by organisms, decomposers take up or excrete nitrogen to maintain their homeostasis (Daufresne & Loreau, 2001). Decomposers limitation determines whether decomposition (*ϕ_D_*, *i.e.,* consumption of detritus flux) is constrained by nitrogen immobilization (*ϕ_I_*; N-limitation) or whether decomposition determines nitrogen dynamics (C-limitation). This link between decomposition and immobilization fluxes, which allows decomposers to maintain their homeostasis is detailed in Appendix S3.

#### Consumer dynamics

Grazers and consumers of decomposers feed on the basal species in their respective ecosystems at a rate *a_G_*, *a_C_* respectively, which follows the law of mass action (*i.e.,* type I functional response): *f_G_* = *a_G_P_CT_ G_CT_* and *f_C_* = *a_C_B_CA_C_CA_* (Appendix S6 details a donor-controlled version of our model). Due to inefficiencies of assimilation, only fractions *e_G_* and *e_C_* of consumption support consumer growth, while the rest is lost from the meta-ecosystem. In addition, to maintain their homeostasis, consumers excrete the excess nitrogen due to the stoichiometric imbalance between their food and their needs (*e.g., β_G_ f_G_* where *β_G_* = (*α_P_ − e_G_α_G_*) for grazers) (Grover & Holt, 1998; Marleau & Guichard, 2019).

#### Nutrient and resource flows

Recycling occurs through the production of detritus of each biotic compartment at rate *d_X_ ∈ {G*, *P*, *C*, *B}*, fueling the detritus pool, and the mineralization of detritus into nitrogen at a rate *m_A_*, in the aquatic system, which corresponds to decomposer’s nitrogen mineralization (Daufresne & Loreau, 2001), and a rate *m_T_* in the terrestrial ecosystem, where the action of decomposers is implicit.

Both ecosystems are supplied with autochthonous nitrogen and detritus flows *I_NZ_* and *I_DZ_* respectively, and detritus and nitrogen are lost at rates *l_NZ_* and *l_DZ_* (*Z ∈* [*T*, *A*]). We assume that inputs of autochthonous detritus do not vary the N:C ratio of detritus.

#### Meta-ecosystem flows

Ecosystems are coupled by spatial flows of subsidies (*e.g.,* detritus from leaching or flooding material decomposition), with a fraction Δ of the produced detritus and excreted nitrogen being directly transferred from each biotic compartment in one ecosystem to the detritus and nitrogen pools in the other ecosystem (regional flow) while the remaining fraction being locally recycled (1 *−* Δ, local flow).

Therefore, we obtain the following set of equations for the terrestrial ecosystem (Fig. 2):

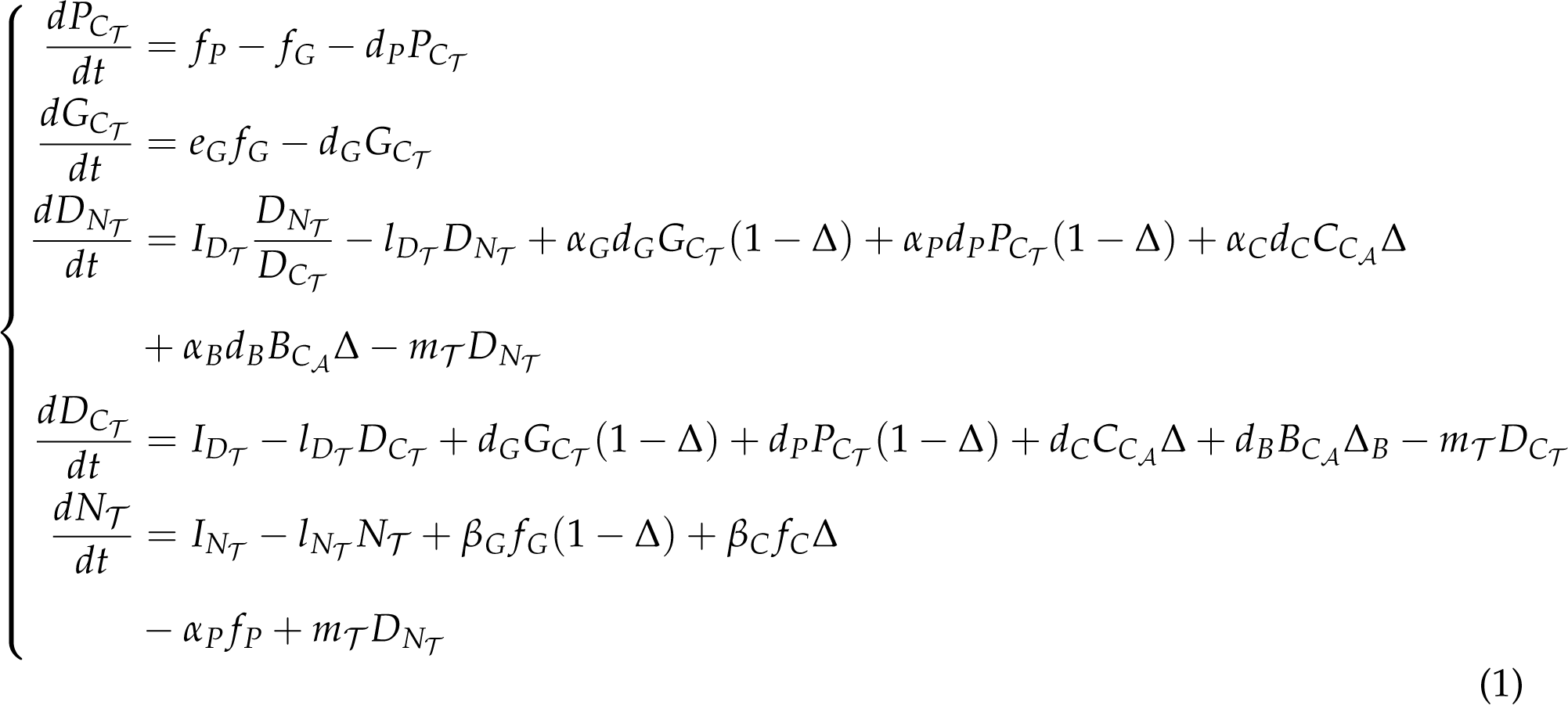

Similarly, we get the following equations for the aquatic ecosystem dynamics:

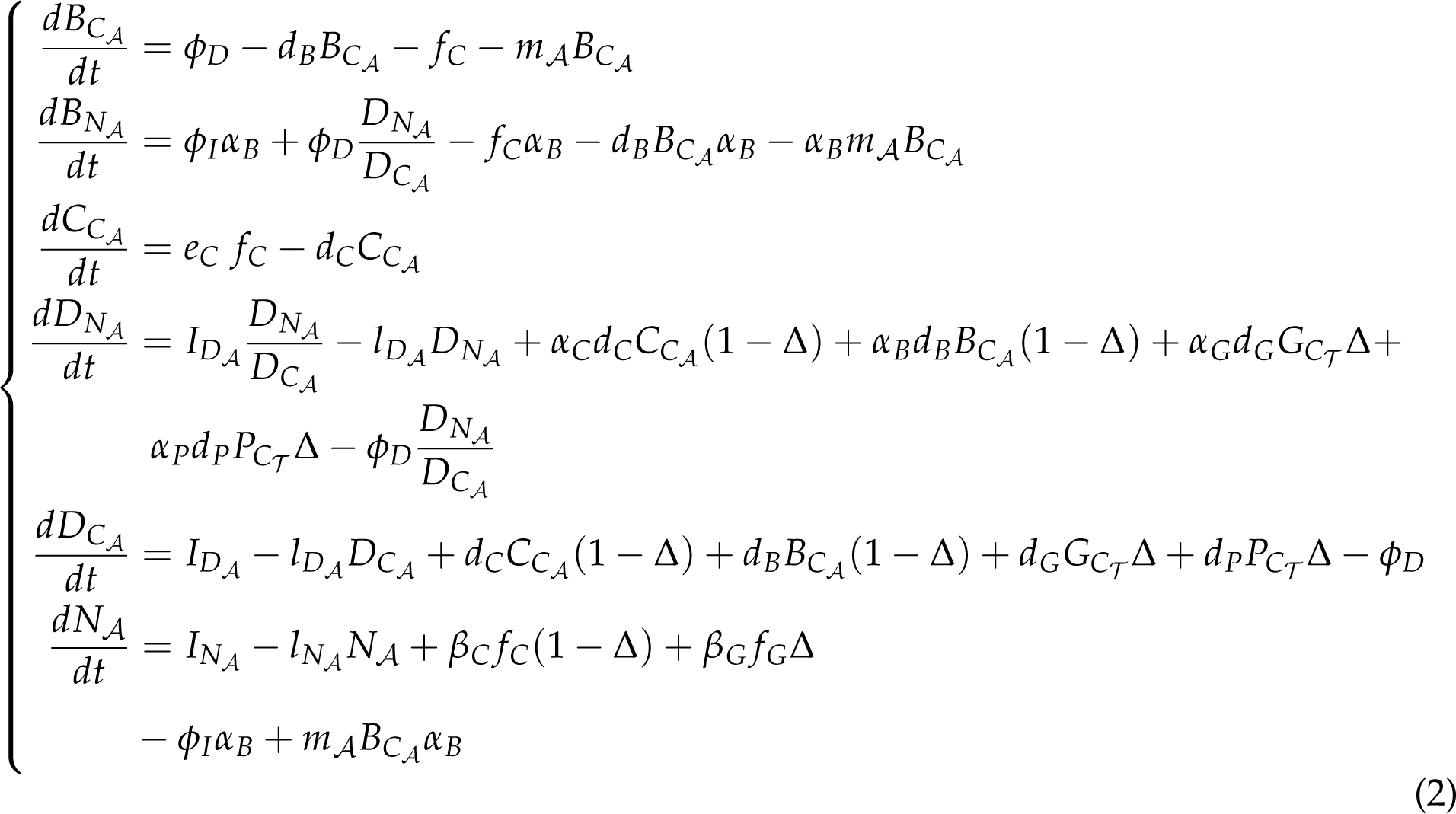

A fully annotated system of equations is available in Appendix S3.

#### Ecosystem & meta-ecosystem metrics

To test our hypotheses on the effect of subsidy stoichiometry on meta-ecosystem functioning, we compared the following ecosystem and meta-ecosystem metrics for two different scenarios of decomposer limitation: nitrogen versus carbon limitation (see parameters in Appendix S3). Under each scenario, we made sure decomposer limitation remained unchanged by the studied variations in decomposer and plant stoichiometry.

#### Production and spatial subsidy flows

We measured the basal and secondary productions (in carbon units) at the ecosystem and meta-ecosystem scales, defined as the flow at equilibrium to basal species (plants and decomposers) and to consumers, respectively.

When decomposer growth is limited by nitrogen, basal production corresponds to immobilization flux of nitrogen (*ϕ_I_*), while it is the decomposition flux (*ϕ_D_*) under carbon limitation. We compared the production of the meta-ecosystem when ecosystems are connected versus isolated (*i.e.,* local recycling of detritus) while varying the stoichiometry of plants and decomposers, using a log ratio response (LRR) defined as:

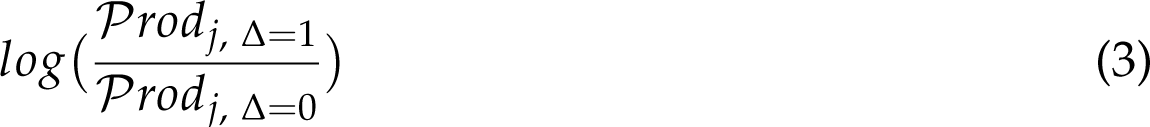

where *Prod_j_*is the production of the trophic level *j* from both ecosystems (meta-ecosystem scale). A positive LRR value means that production benefits from subsidy flows relative to a scenario where all subsidies are locally recycled (no spatial flows).

#### Measuring the feedback between ecosystems

We measured the spatial effects (hereafter feedbacks) of subsidy flows as the difference in production between scenarios where ecosystems are bidirectionally versus unidirectionally connected by subsidy flows. In the latter, the focal ecosystem receives subsidies from the connected ecosystem but its own exported flows are lost from the meta-ecosystem. For instance, for the case of the terrestrial ecosystem, we aim to quantify how resources it exports impact its own production through the effect of spatial flows exported to the aquatic ecosystem and the quantity of aquatic subsidies exported back (see Fig. S1.1). We interpret this measure as the strength of the spatial feedback *F* induced by the subsidy flow:

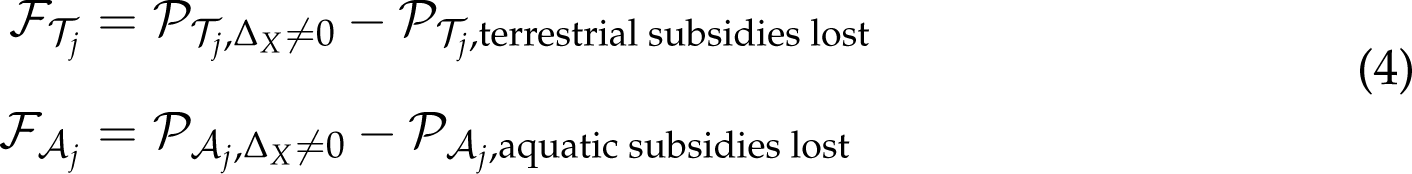

where *P_Z__j_* is the production of the trophic level *j*, *j ∈* [1, 2] in ecosystem *Z*, *Z ∈* [*T*, *A*]. In addition, because this metric evaluates the long-term impact of spatial flows on each ecosystem production, we also measured the short-term response of each trophic level density to a transient increase in the fraction of subsidies received by the other ecosystem (Appendix S1).

#### Parameters and sensitivity analysis

We parameterized the model to account for differences of functioning between aquatic and terrestrial ecosystems, and of stoichiometry between plants and decomposers, while staying generic and covering the different qualitative behaviors that might be observed in different systems (see Appendix S3, and Table S3.3). In addition to the sensitivity analyses on the shape of the functional responses, and the number of trophic levels, we performed one on the parameter values (Appendices S6-S7). As the model is not analytically tractable, we performed simulations using *DifferentialEquation* package in Julia (version 1.7.3), and analyzed them in R (version 4.1.0). Details about the simulation method are given in Appendix S3.

## Results

The following results hold qualitatively when changing the type of functional response, adding a third trophic level, modifying the asymmetry of flows between ecosystems, and varying the parameter values (Appendices S4, S6-8).

### Spatial complementarity under carbon limitation

Under the scenario of carbon limitation for aquatic decomposers, resource spatial flows maximize both primary and secondary production in each local ecosystem when decomposers have high nitrogen content (*α_B_*) and plants have low nitrogen content (*α_P_*) (Fig. 3A). In addition, spatial flows also increase the densities of grazers and aquatic consumers (Fig. S1.2).

**Figure 3:**
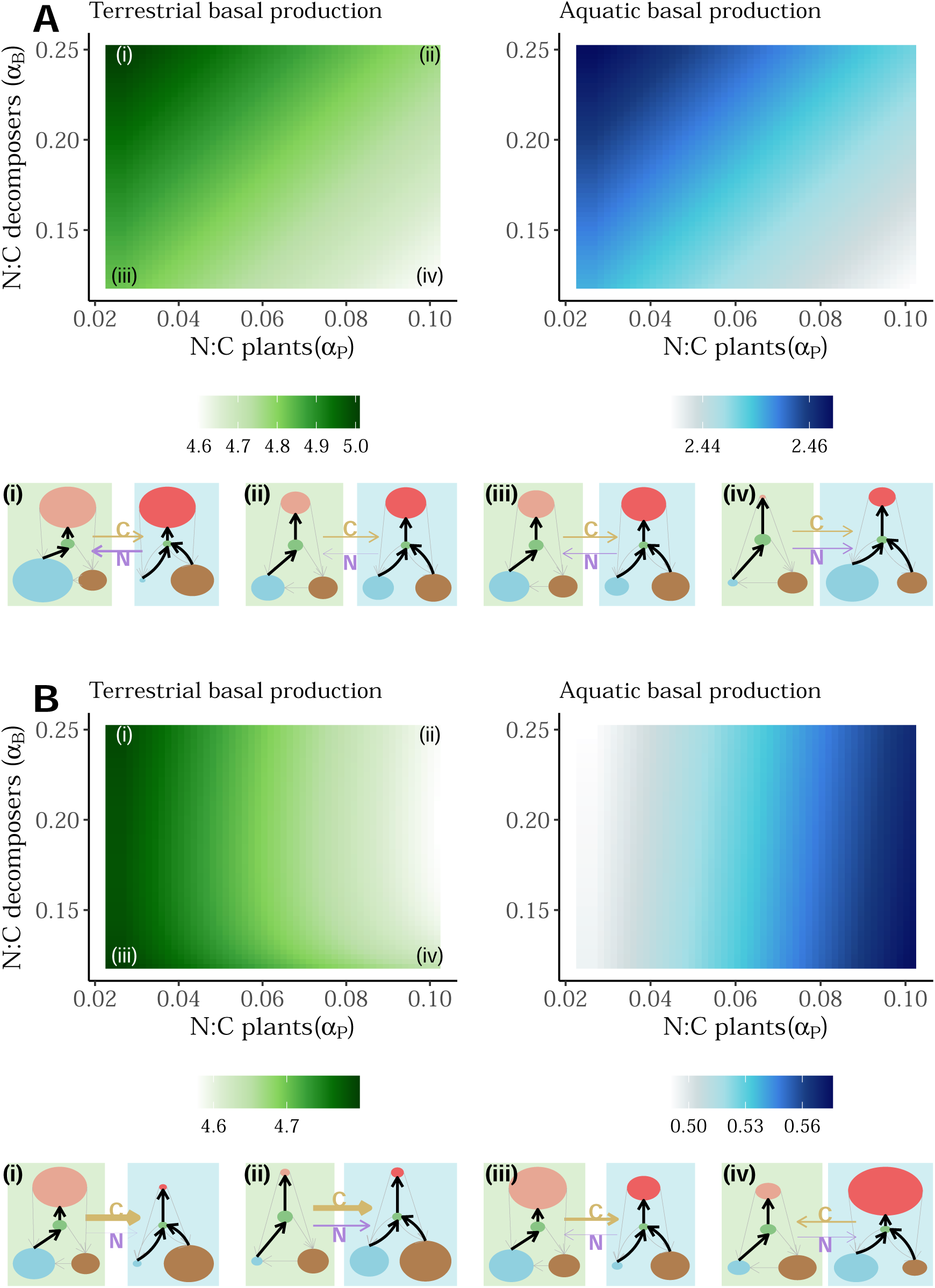
Effects of basal species limitation and stoichiometry on ecosystem production. We show the basal production in terrestrial (left) and freshwater (right) ecosystems with spatial coupling (Δ = 1, see Fig. S7.5 in appendix S5 for varying values) according to the N:C ratio of basal species. In (A), decomposers are carbon limited, while they are limited by nitrogen in (B). Production in both ecosystems is expressed in carbon units. The results for secondary production are depicted in Fig. S1.6. We also represent the meta-ecosystem compartments at equilibrium when ecosystems are connected Δ = 1 (terrestrial ecosystem in green and freshwater one in blue) for the four corners of the heat map (*(i)*: *α_B_* = 0.25, *α_P_* = 0.025, *(ii)*: *α_B_* = 0.25, *α_P_* = 0.1, *(iii)*: *α_B_* = 0.12, *α_P_* = 0.025, *(iv)*: *α_B_* = 0.12, *α_P_* = 0.1). Black arrows correspond to trophic interaction while gray arrows correspond to recycling paths. The two trophic levels are represented in pink for primary consumers and green for basal species. Nitrogen and detritus pools are respectively in blue and brown. For detritus, we only represent the carbon content. The size of each compartment is set relative to the other same compartment to allow direct size comparison across the stoichiometric conditions. We also represent the net flow of carbon (orange arrow) and nitrogen (purple arrow) at the meta-ecosystem scale. Their width is proportional to the flow.

As decomposers need more nitrogen to maintain their homeostasis (*α_B_* increases), both the nitrogen content of decomposer-produced detritus and the quantity of nitrogen excreted by aquatic consumers through stoichiometric imbalance (*β_C_* = *α_B_ − e_C_α_C_*) increase. When ecosystems are connected, this flow of nitrogen fuels the terrestrial nitrogen pool, and supports higher plant production (Fig. 3A-*(iii) to (i)*) and higher density of grazers (Fig. S1.2). These mechanisms contribute to higher carbon and nitrogen flows exported back to the aquatic ecosystem. In addition, we observe that the aquatic ecosystem is a net source for nitrogen and a net sink for carbon at the meta-ecosystem scale (Fig. 3A *(i)-(iii)*). Production in both ecosystems is maximized for plants with a low N:C ratio (low *α_P_*, Fig. 3A). When plant N:C decreases (*e.g.,* from Fig. 3A-*(ii) to (i)*), basal plant production increases due to lower needs of nitrogen for the same growth, and consequently leads to a higher grazer density (Figs S1.2, 3A-*(ii)*). Thus, the quantity of detritus produced by plants and grazers, and then exported, increases (Fig. S1.3). Consequently, it supports higher production in the aquatic ecosystem (Fig. S1.2). Additionally, because resources exported from the terrestrial ecosystem have on average a lower N:C ratio compared to the ones produced by the aquatic ecosystem, the N:C ratio of aquatic detritus decreases (Fig. S1.4B), which relaxes the decomposer limitation for carbon (Fig. S1.4C) and could, in specific conditions, induce a change in the limitation of decomposers (from carbon to nitrogen, Fig. S1.5).

Besides, at the meta-ecosystem scale, spatial coupling between ecosystems mainly lead to higher primary and secondary production than the sum of the production of isolated systems (Fig. 4A). Positive effects result from a stoichiometric matching between the local community needs and the spatial flows (*e.g,* plants need nitrogen that is brought by nitrogen-rich aquatic subsidies), suggesting a spatial complementarity of ecosystems.

**Figure 4:**
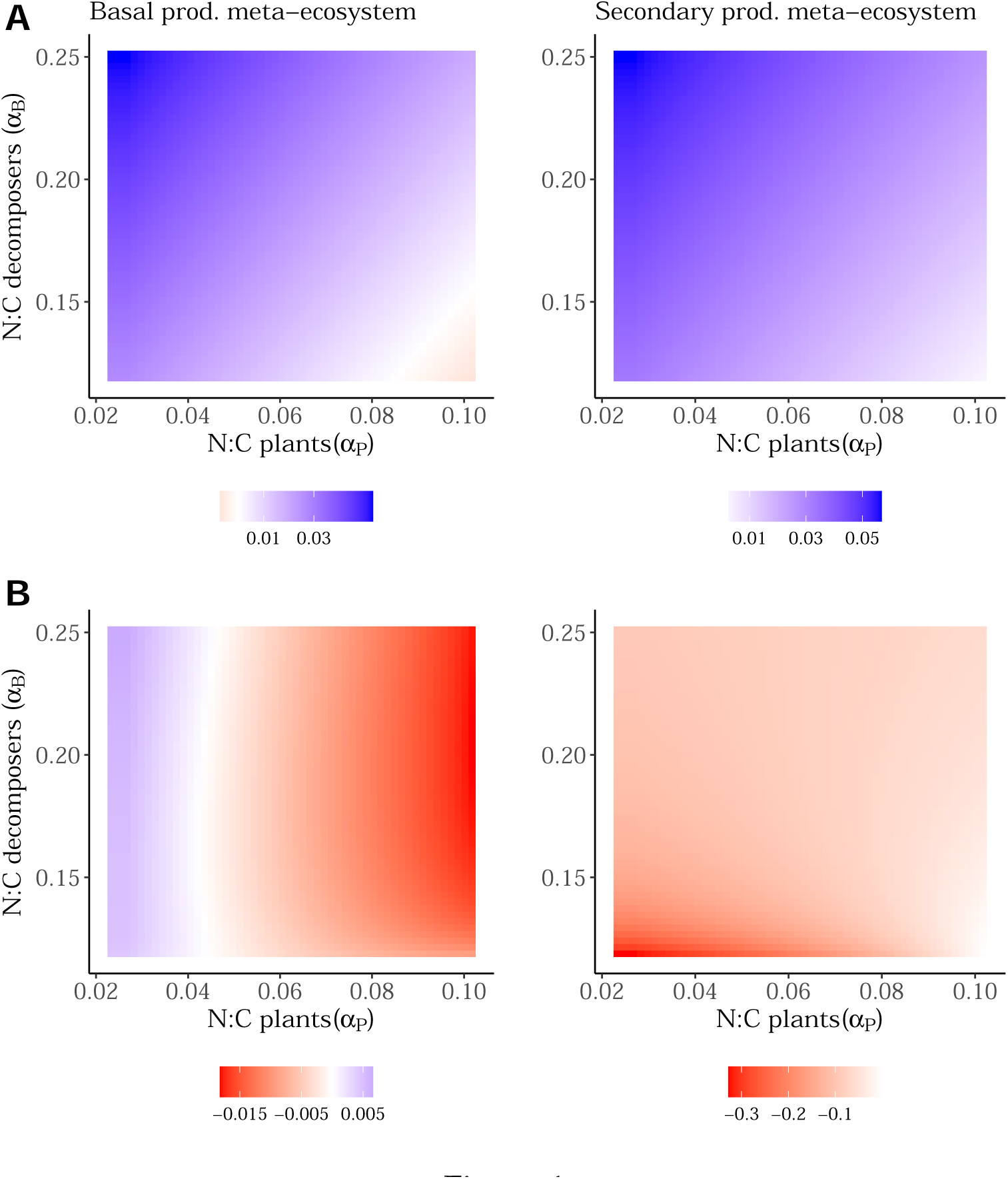
Effect of spatial flows on meta-ecosystem production. We compare the log ratio-response of basal production (plants + decomposers) in left panels, and secondary production (grazers + consumers of decomposers) in right panels, at the meta-ecosystem scale when ecosystems are connected versus isolated. (A): Decomposers are carbon limited. (B) Decomposers are nitrogen limited. The black line indicates the isocline delimiting positive and negative effects of spatial subsidies. We provide in Fig. S1.8 a scheme that summarizes the mechanism.

### Ecosystem spatial competition under nitrogen limitation

Conversely, under nitrogen limitation of decomposers, we observe opposite patterns between ecosystems: spatial flows maximize production in one ecosystem when minimizing it in the other. Basal production in the aquatic ecosystem peaks for high *α_P_* (*i.e.,* nitrogen-rich plants), when plants produce nitrogen-rich detritus, while it peaks for low *α_P_* in the terrestrial ecosystem, when plant demand for nitrogen is low (Fig. 3B). When *α_B_* increases, it exacerbates the stoichiometric mismatches between decomposers and both their consumers and their detrital resources (*i.e.*, 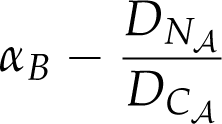 increases; Fig. S1.4). The stoichiometric mismatch with consumers reduces secondary production (Fig. S1.2B). The stoichiometric mismatch with detritus reduces the nitrogen pool in the aquatic ecosystem (because decomposers need more nitrogen to maintain their homeostasis) and limits the decomposition of detritus (Fig. 3B). In these conditions (high *α_B_*), the aquatic ecosystem acts as a net sink receiving on average more carbon and nitrogen flows than it exports (Fig. 3B). We find the same pattern in empirical data on terrestrial-freshwater exchanges (Fig. 1C; Box 1). Additionally, we observe that meta-ecosystem primary and secondary production generally decrease when both ecosystems are connected (Fig. 4B), which denotes a spatial competition rather than complementarity effect between ecosystems: the resources exported by the terrestrial ecosystem do not match the aquatic community needs.

### Cross-ecosystem feedbacks

Lastly, we measured how the subsidies exported by a donor ecosystem, feedback on its production through the subsidies exported by the recipient (Figs. 5, S1.1). Under carbon limitation of decomposers, feedbacks on basal and secondary production in both ecosystems are always positive and scale non-linearly with the fraction of subsidies regionally exported (Figs. 5A, S1.7a). This means that exports of resources from the donor ecosystem, compared to when being lost, always increase production of the recipient ecosystem, which further increases the fraction of subsidies exported back. The strength of this positive feedback is highly and positively modulated by the stoichiometry of decomposers, while plant stoichiometry only slightly changes the feedback (Fig. 5A). Conversely, feedbacks reach negative values under nitrogen limitation of decomposers, meaning that production is lower with bidirectional flows than with a unidirectional flow leaving the meta-ecosystem (Figs 5B, S1.7b). Indeed, we found a U-shape relationship crossing 0 between the fraction of subsidies being transferred regionally and the strength of the feedback in the terrestrial ecosystem.

**Figure 5:**
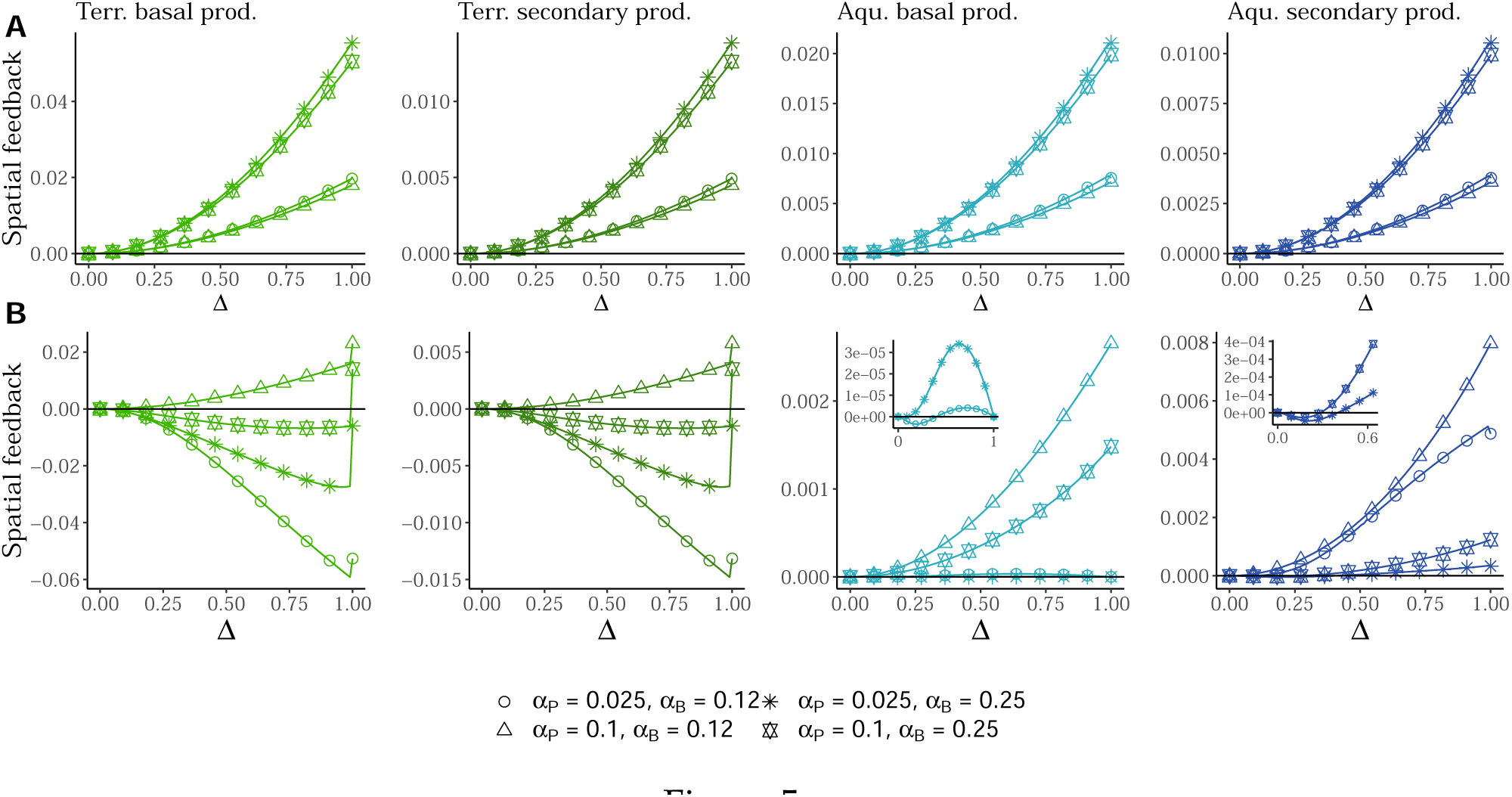
Feedback between ecosystems depends both on decomposer limitation and basal species stoichiometry. In order to understand how resources exported by an ecosystem feedback on its own production through the effect on the recipient ecosystem, we compared for each trophic level and both ecosystems the production of the recipient ecosystem for scenarios of bidirectional and unidirectional flows of subsidies along a gradient of connectivity between ecosystems (Δ; see methods and Fig. S1.1 for formulas). Shapes correspond to the four extreme scenarios of basal species stoichiometry defined in Fig. 3. of plants and decomposers (see Fig. 3). (A): C-limited decomposers and (B): N-limited decomposers. Some inserts are displayed for some curves to zoom in and better visualise the interplay between stoichiometric mismatch and mass-effect mechanisms (*i.e.*, the sign of the feedback).

At first, the feedback decreases with increasing connectedness of ecosystems (for Δ *∈* [0, 0.25] and Δ *∈* [0, 0.9] in aquatic and terrestrial ecosystems, respectively) due to an amplification of the stoichiometric mismatch between decomposers and their detritus following carbon-rich terrestrial subsidies inputs. However, at higher regional flows, the terrestrial ecosystem exports more detritus regionally due to higher levels of nitrogen-rich subsidies from the aquatic ecosystem. The mass effect of abundant terrestrial subsidies overcomes the negative saturating effect of stoichiometric mismatch between detritus and decomposers (Fig. S1.7c), thus allowing the terrestrial system to sustain higher aquatic production compared to a scenario where subsidies from the aquatic ecosystem are lost. We summarized the mechanisms driving spatial feedbacks at terrestrial-aquatic ecotones in Fig. S1.8.

## Discussion

By analyzing a stoichiometrically explicit meta-ecosystem model, we reveal how local ecosystem limitations and basal species stoichiometry drive meta-ecosystem functioning, by leading either to stoichiometric complementarity or competition between spatial flows and local demand. Our results highlight the potential for complementarity in resource use among ecosystems at the landscape scale, but also show that resource flows can induce negative spatial feedback between terrestrial and freshwater ecosystems, depending on local nutrient limitations. We discuss the implications of these findings and provide perspectives for spatial ecology.

### Spatial complementarity vs competition through subsidy flows

Under carbon limitation of aquatic decomposers, spatial flows maximize production in both ecosystems for basal species with complementary needs (low-N plants and high-N decomposers). Terrestrial subsidies alleviate carbon limitation and sustain production in the aquatic ecosystem, while nitrogen-rich subsidies from the aquatic ecosystem allow plants of the terrestrial ecosystem to fix more carbon, together leading to higher meta-ecosystem production. Both recent experimental works and reviews have stressed the importance of spatial flow quality for local ecosystem functioning (Sitters <u>et al</u>., 2015; Gounand <u>et al</u>., 2017). Using two-patch microcosms, Gounand et al. showed that detritus exchange between autotrophic and heterotrophic communities can maximize densities in both communities compared to isolated ones (Gounand <u>et al</u>., 2017). In a recent meta-ecosystem modeling work, (Harvey <u>et al</u>., 2021) propose the cross-ecosystem efficiency hypothesis to describe how spatial flows might foster higher functions at the landscape scale by redistributing resources among ecosystems differing in the functions they optimize (primary production in terrestrial ecosystems versus higher trophic efficiency in aquatic ecosystems). We further propose that complementarity between ecosystem can arise from the match between the stoichiometry of spatial flows and local community needs, leading to higher production when ecosystems are connected. Indeed, the complementarity of ecosystems creates a spatial positive feedback loop: on the one hand, autotrophic ecosystems, export carbon-rich organic matter that fuels heterotrophic ecosystems and fosters local production. In return, more nutrient-rich subsidies are exported back, promoting primary production and closing the meta-ecosystem loop. Therefore, we propose that cross-ecosystem flows bound ecosystems into a spatial auto-catalytic loop (*sensu* Veldhuis <u>et al</u>., 2018), maximizing the production of the whole.

However, cross-ecosystem effects are highly modulated by local ecosystem limitations. While we find that connectedness leads to complementarity when the aquatic ecosystem is limited by carbon, a mismatch arises between aquatic community needs and resources exported by the terrestrial ecosystem under nitrogen limitation. In this case, plants and decomposers indirectly compete at the meta-ecosystem scale for nitrogen: high nitrogen demand and low-quality terrestrial subsidies lower decomposition and production in the aquatic ecosystem, leading to negative spatial feedback on terrestrial primary production. Then, both consumers of decomposers and grazers shrink in densities due to reduced basal production. In non-spatial experiments, decomposers often evolve toward co-limitation as a mechanism to avoid competitive exclusion when competing for nutrients with autotrophs (Daufresne & Loreau, 2001; Danger <u>et al</u>., 2008; Daufresne <u>et al</u>., 2008). In our meta-ecosystem model, N-C co-limitations in decomposers always trigger positive spatial feedbacks (see Appendix S5), suggesting that if such evolution was to occur in a spatial context, it would maximize production at the meta-ecosystem scale.

### Understanding cross-ecosystem interactions through resource feedbacks

Cross-ecosystem facilitation (*i.e.,* complementarity mechanism) and competition (*i.e.,* mismatch mechanism) occur through spatial feedback, which sign is modulated by the limitation of decomposers and the stoichiometry of basal species. In particular, we emphasize that feedbacks on terrestrial production could reach negative values under nitrogen limitation of the aquatic ecosystem while they are always positive and scale non-linearly with the connectedness of ecosystems under carbon limitation. In addition, feedbacks on the production of both ecosystems increase with the N:C ratio of decomposers. While our model is spatial, recent models connecting brown and green food chains in non-spatial contexts also emphasized the importance of microbial stoichiometry in determining the interaction between decomposers and plants (Zou <u>et al</u>., 2016; Buchkowski <u>et al</u>., 2019). Zou <u>et al</u>. (2016) showed how the impact of the green food web on the brown one was highly determined by their resource limitation of decomposers: the effect was either positive or negative when decomposers were C- or N-limited respectively. In a meta-ecosystem perspective, we found that feedbacks between autotrophic and heterotrophic food-webs reach negative values under nitrogen limitation: spatial flows of terrestrial subsidies negatively impact aquatic ecosystem production through a direct stoichiometric mismatch effect, and may indirectly decrease terrestrial ecosystem production compared to a scenario where terrestrial resources exported are lost from the system. This finding questions the term ”subsidies” (*sensu* Polis <u>et al</u>., 1997; but see Subalusky & Post, 2019), defined as a positive effect on production in the recipient ecosystem. This stoichiometric mechanism contrasts with previous theoretical works, which suggested that cross-ecosystem flows always increase landscape production (*e.g.,* Marleau <u>et al</u>., 2014). In fact, stoichiometric mismatch is an important constraint upon consumer growth (Hillebrand <u>et al</u>., 2009). In an experimental context, several authors stressed that lower food quality can decrease consumer assimilation efficiency and population growth rates, and further lead to population decrease in aquatic systems (*e.g.,* Anderson <u>et al</u>., 2005; Brett <u>et al</u>., 2009; Hillebrand <u>et al</u>., 2009). For instance, Kelly et al. (2014) showed a negative association between zooplankton production and inputs of terrestrial organic carbon in lakes caused by shading and low quality of terrestrial inflows (Kelly <u>et al</u>., 2014).

By contrast, Pacific salmons that annually migrate to freshwater ecosystems can relax nitrogen limitation of heterotrophic bacteria, leading to higher aquatic production (Rü egg <u>et al</u>., 2011). Salmons can further be eaten by bears, increasing both their excretion rates and the carcass deposition and contributing to a nitrogen flow positively affecting riparian forests (Helfield & Naiman, 2006 ; Box 1). Besides, empirical studies support our results that cross-ecosystem material flows can shape the functioning within recipient ecosystems by modulating nutrient limitation (see Sitters <u>et al</u>., 2015 for a review). Thus, empirical observations support our results that cross-ecosystem material flows can shape the production of recipient ecosystems by relaxing or exacerbating nutrient limitation (Montagano <u>et al</u>., 2018).

### Relaxing some model assumptions

As for any modelling studies, we derived mechanisms and predictions at the cost of simplifying hypotheses on processes. First, we assumed that each organism was held at a constant stoichiometry. Recent experiments or meta-analyses suggest that decomposers and their consumers can deviate from strict homeostasis (Cross <u>et al</u>., 2003; Persson <u>et al</u>., 2010; Scott <u>et al</u>., 2012) Similarly, plant stoichiometry often changes depending on the dominance of species and their traits (*e.g.* Yu <u>et al</u>., 2010), and nutrient inputs into terrestrial ecosystems. We would expect this plasticity mechanism to reduce the stoichiometric mismatch between terrestrial exports and aquatic needs as the nitrogen content of plants would increase in response to nitrogen-rich aquatic subsidies and the decomposers would have reduced N:C ratio following carbon-rich terrestrial inputs. In the example of salmon, nitrogen actively transferred to terrestrial ecosystems by bears increases the N:C ratio of plant foliage (Hilderbrand <u>et al</u>., 1999). Additionally, we ignored compensatory feeding mechanisms that consumers can develop when facing low-quality food (Cebrian & Lartigue, 2004; Hillebrand <u>et al</u>., 2009). Yet, Yet, these mechanisms can promote extinction of producers through consumer overfeeding (Marx <u>et al</u>., 2021) and decrease trophic efficiency of terrestrial and aquatic ecosystems (Frost <u>et al</u>., 2006; Hillebrand <u>et al</u>., 2009). In addition, at the ecosystem level, a balance between top-down and bottom-up control also emerges depending on whether subsidies are direct resources or not (as reviewed by Allen & Wesner, 2016). For instance, using models (McCary <u>et al</u>., 2021) and (Leroux & Loreau, 2008) showed that the net control induced by subsidy flow depends on the trophic position of the species consuming the subsidy in the recipient ecosystem as well as its preference for this subsidy. Accounting for both top-down and bottom-up effects at both physiological (compensatory feeding and deviation from strict homeostasis) and ecosystem level (net control) may help to better predict community response to spatial subsidies.

Secondly, the impact of subsidy quality can be explored using a different framework than ecological stoichiometry such as nutritional ecology (Raubenheimer <u>et al</u>., 2009; Schindler & Smits, 2017; Osakpolor <u>et al</u>., 2023). At the terrestrial-aquatic ecotone, if carbon-rich subsidies exported by the terrestrial ecosystem decrease decomposer growth-efficiency due to carbon being recalcitrant, the strength of the positive feedback emerging at the landscape level would decrease (Fig. S7.4).

### Perspectives at the landscape scale

We stress that future models should go beyond spatially implicit models to account for landscape structure and its heterogeneity. A promising future path would be to link the spatial distribution of resources and foraging behavior of consumers. Consumers often avoid areas with low-quality resources (*e.g.,* Duparc <u>et al</u>., 2020; Rizzuto <u>et al</u>., 2021). By doing so, they limit important stoichiometric mismatch with their food and avoid nutrient limitations. Modelling studies integrating such processes would help for instance understanding how animal foraging movements interact with predation avoidance to determine the distribution of nutrients at the landscape scale (Anderson <u>et al</u>., 2010; Schmitz <u>et al</u>., 2018; Ferraro <u>et al</u>., 2022). This may highlight further emergent patterns such as nutrient co-limitation induced by spatial processes (Marleau <u>et al</u>., 2015) or patchy distribution of resources (Johnson-Bice <u>et al</u>., 2022). Our study sets bases to understand fundamental mechanisms by which spatial flow stoichiometry modulates cross-ecosystem interactions and the functioning of communities at different scales. Integrating perspectives of movement ecology and ecological stoichiometry should help building a more integrative spatial ecology better accounting for the interplay between biogeochemical cycles and community dynamics at the landscape extent.

#### Box 1

Empirical cross-ecosystem subsidy flows between freshwater and terrestrial ecosystems

We gathered data from the literature on cross-ecosystem flows between freshwater (streams, lakes) and terrestrial (forests, grasslands) ecosystems and provide here orders of magnitudes and stoichiometric characteristics of these fluxes (see Appendix S1 for details on data extraction). First, at the freshwater-terrestrial ecotone, subsidies are mostly represented by plant litter exported by terrestrial ecosystem and invertebrates exported by both ecosystems (Fig. 1A). Interestingly, exported materials by terrestrial *versus* freshwater ecosystems vary in quantity (Fig. 1C) and quality (carbon and nitrogen content; (Fig. 1B, Table S2.2)). While the exports of terrestrial ecosystems, mainly characterized by plant litter, are of poor quality (*i.e.*, carbon-rich), resources from freshwater ecosystems are richer in nitrogen. Aside from quality, these cross-ecosystem flows also vary in quantity. When comparing the carbon and nitrogen flows at terrestrial-freshwater ecotone, we observe a strong asymmetry of carbon flows crossing ecotones (20 *gC*.*m*^−2^.*yr*^1^ exported in average by freshwater ecosystems *versus* 128 *gC*.*m*^−2^.*yr*^1^ from terrestrial ones; Fig. 1C & Table S2.3). However, this quantitative asymmetry of exported resources is partially compensated by their stoichiometric asymmetry (see Appendix S8). Consequently, while there is one order of magnitude of difference for carbon flows, we only observe a slight difference for nitrogen flows.

Thus, we observe two types of asymmetries in resource flows at freshwater-terrestrial ecotone. First, there is an asymmetry in flow quantity that may originate from differences in convexity of ecosystem profiles (*e.g.* forest *versus* stream ; see Polis <u>et al</u>., 1997; Shurin <u>et al</u>., 2006; Leroux & Loreau, 2008). Second, there is an asymmetry in quality, which originates from stoichiometric differences of organisms (Elser <u>et al</u>., 2000) and can impact local ecosystem functioning. In fact, high quantity but low quality of terrestrial subsidies can limit freshwater ecosystem production by various mechanisms ranging from shading of lakes, increase in stoichiometric mismatch or reduction of freshwater production (*e.g.,* (Kelly <u>et al</u>., 2014)). Similarly, high quality subsidies can considerably impact terrestrial plant communities in both richness and production (Helfield & Naiman, 2006; Rü egg <u>et al</u>., 2011; Hocking & Reynolds, 2011; Bultman <u>et al</u>., 2014). Together, it emphasizes the need to move toward a stoichiometric theory at the landscape extent (Leroux <u>et al</u>., 2017).

## Author contribution

All authors conceived the study. B.P performed research and wrote the first draft, which was substantially revised by E.T, G.L and I.G.

All authors read and approved the final manuscript.

## Acknowledgement

We thank Frédéric Guichard for interesting discussions and feedbacks on a previous version. We are grateful to the three anonymous reviewers for their thorough assessment of our work and for providing constructive and insightful comments.

