## Supplementary file for "Quality matters: stoichiometry of resources modulates spatial feedbacks in aquatic-terrestrial meta-ecosystems"

### Contents

**S1 Additional figures referenced in the main text** **S2**

**S2 Extracting cross-ecosystem data flows** **S13**

**S3 Derivation of $\phi_I$ and $\phi_D$, full model system and parameters values** **S16**

**S4 Adding a trophic level** **S26**

**S5 Co-limitation of decomposers** **S31**

**S6 Donor-Control functional responses** **S33**

**S7 Sensitivity analysis on the parameter values** **S37**

**S8 Sensitivity analysis on the asymmetry of flows** **S44**

### S1 Additional figures referenced in the main text

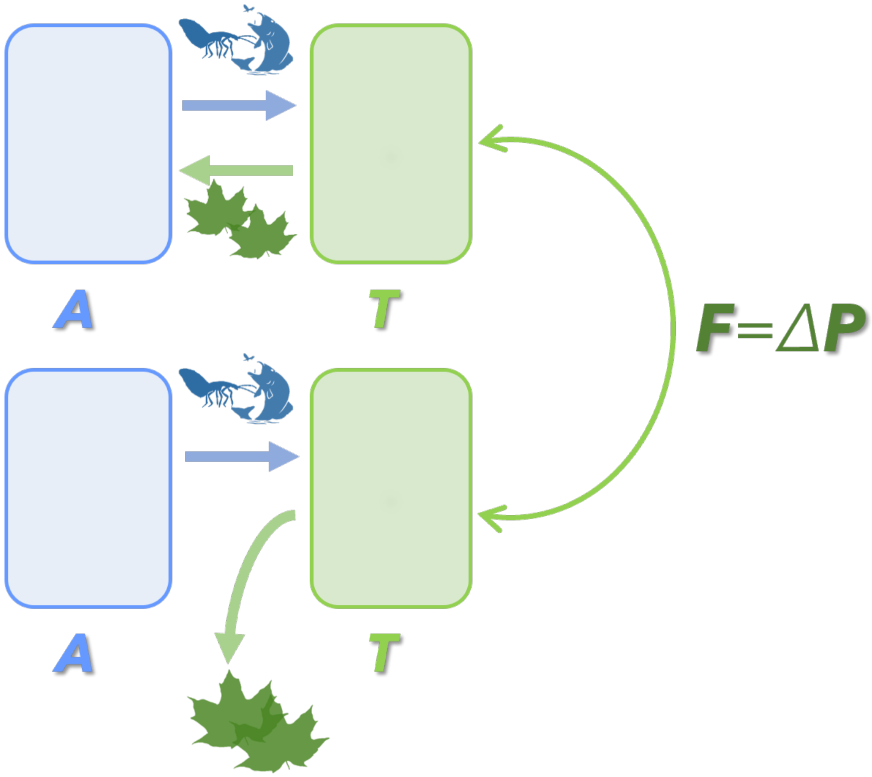


Figure S1.1: **Quantifying the feedback.**
We quantified the feedback ($\mathcal{F}$, here on terrestrial production) by subtracting the terrestrial production ($\mathcal{P}$, either primary or secondary) when ecosystems were bidirectionally connected (top) and when subsidies exported by terrestrial ecosystems were lost from the meta-ecosystem (bottom). Therefore, the feedback strength is defined as a difference in ecosystem functioning between two scenarios: when there is a bidirectional exchange of subsidies or not.

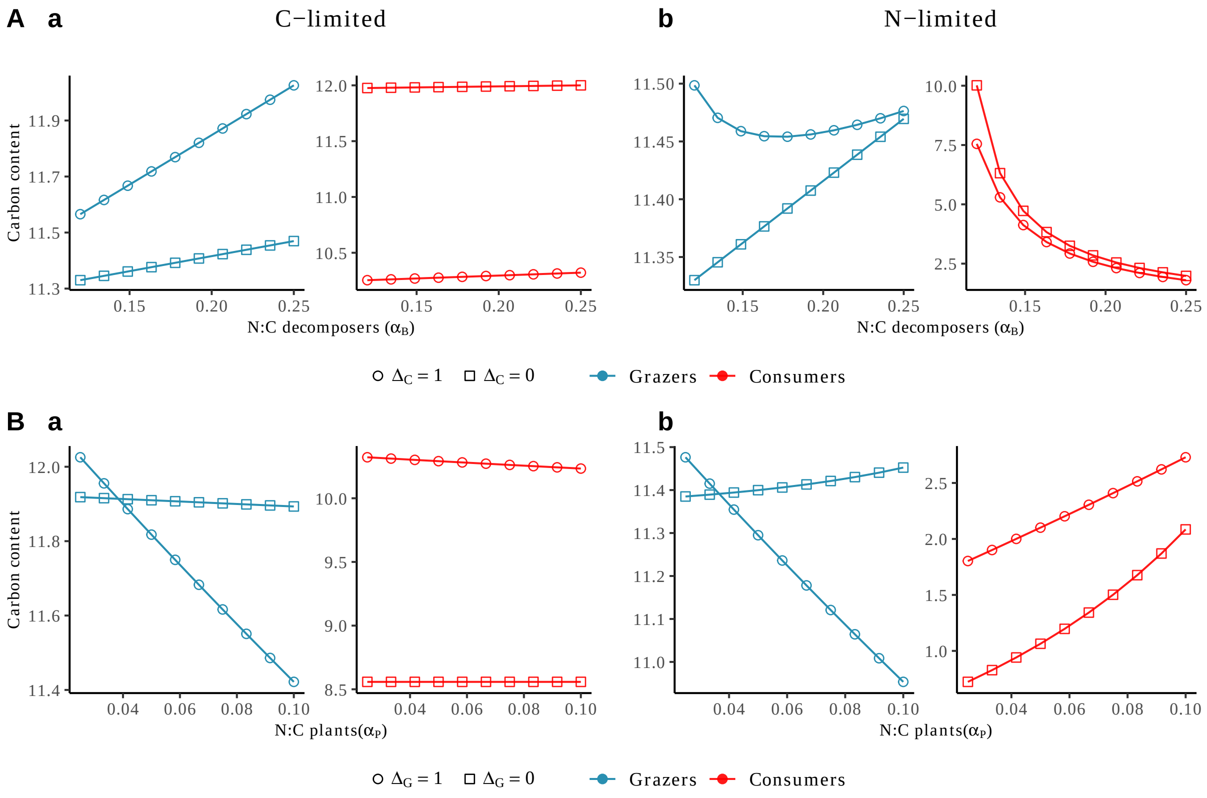


Figure S1.2: **Variations of consumer densities measured by carbon content along the gradients of the stoichiometry of basal species.**
We compare the carbon content of both consumers (red) and grazers (blue) along the gradient of N:C of decomposers ($\alpha_B$ ; A) and plants ($\alpha_P$ ; B) between scenarios where primary consumers subsidies are regionally transported (circles) or locally recycled (squares). Decomposers are either carbon-limited (left; a) or nitrogen-limited (right; b). Other parameters: in (A): $\alpha_P = 0.025$ and $\Delta_C = 0$ meaning that all detritus produced by consumers of decomposers were locally recycled and in (B): $\alpha_B = 0.25$ and $\Delta_G = 0$ meaning that all detritus produced by grazers were locally recycled.

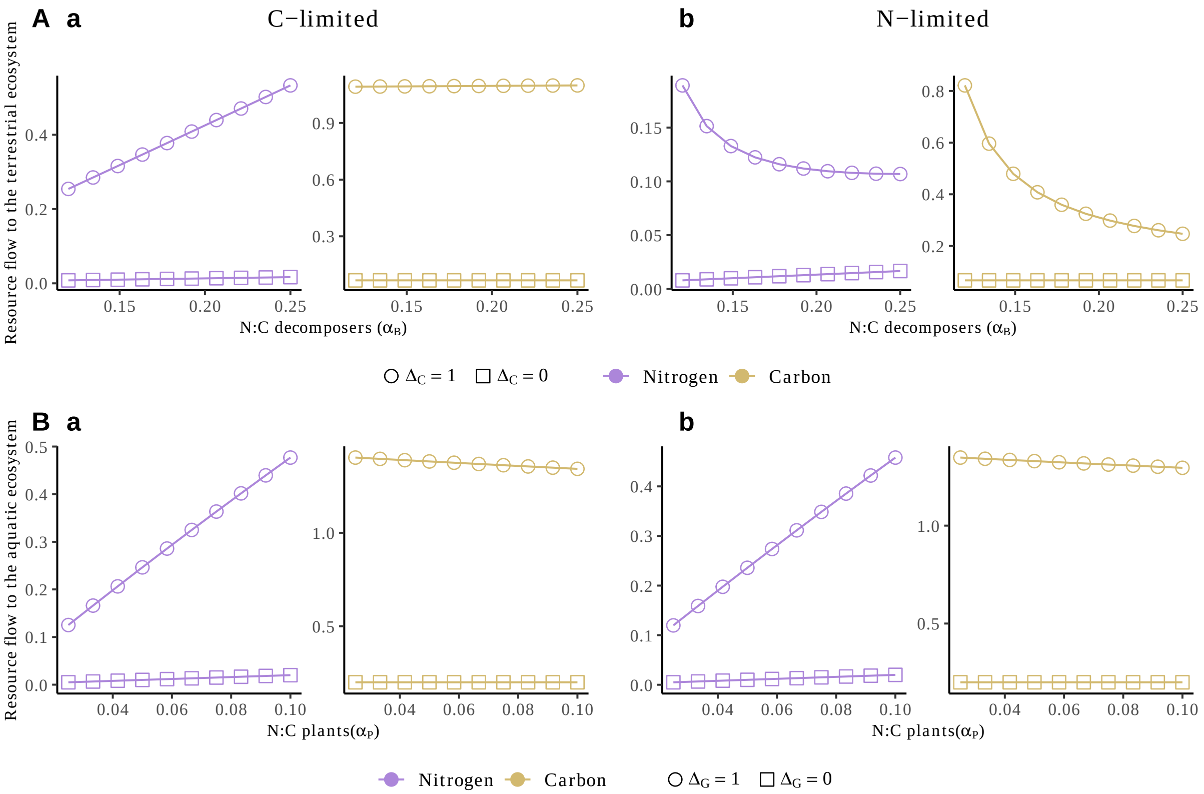


Figure S1.3: **Changes in the flows of carbon and nitrogen exported along the gradients of the stoichiometry of primary producers.**
We compare the flows of resources (carbon in orange, nitrogen in purple) to the aquatic and terrestrial ecosystems along the gradient of N:C of decomposers ($\alpha_B$ ; A) and plants respectively ($\alpha_P$ ; B) between scenarios where primary consumers subsidies are regionally transported (circles) or locally recycled (squares). Decomposers are either carbon-limited (left, a) or nitrogen-limited (right; b). Other parameters: in (A): $\alpha_P = 0.025$ and $\Delta_C = 0$ meaning that all detritus produced by consumers of decomposers were locally recycled and in (B): $\alpha_B = 0.25$ and $\Delta_G = 0$ meaning that all detritus produced by grazers were locally recycled.

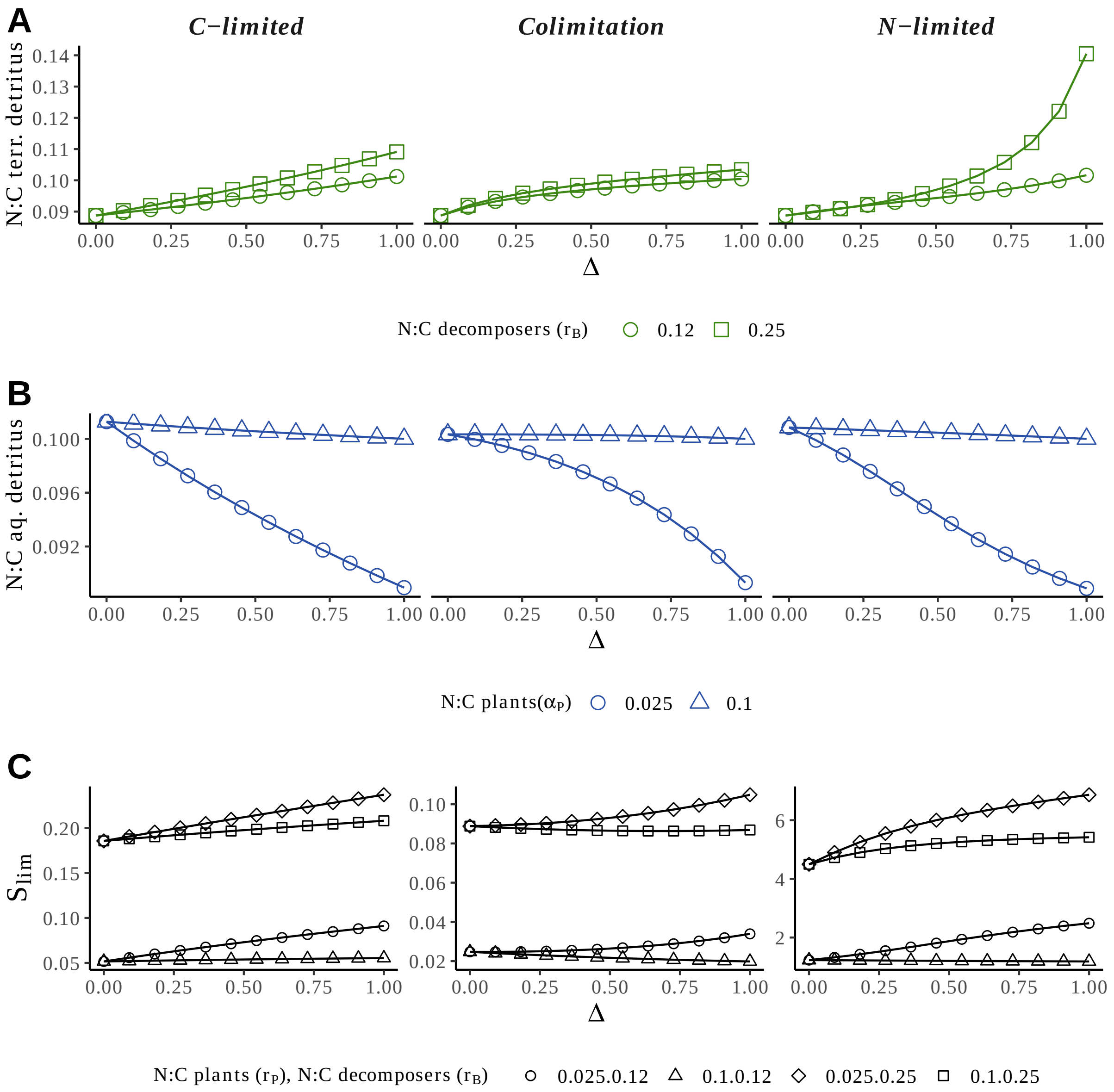


Figure S1.4: **Variation of detritus N:C and stoichiometric ratio $S_{\mathbf{lim}}$ in aquatic and terrestrial ecosystems under the three decomposers limitation**.

We show the variation of the N:C ratio of detritus in both ecosystems (A for Terrestrial (terr.), B for aquatic (aq.)) as a function of the fraction of subsidies being regionally transferred (Δ). We also represent the variation of the stoichiometric ratio $S_{\text{lim}}$ (see Appendix A). A value below (resp. above) 1 indicates that decomposers are under carbon (resp. nitrogen) limitation. These analyses were performed for the three types of limitations of decomposers: C-limitation (left column), co-limitation (middle column), and N-limitation (right column). Other parameters : for (A), $\alpha_P$=0.025 ; for (B), $\alpha_B$=0.12.

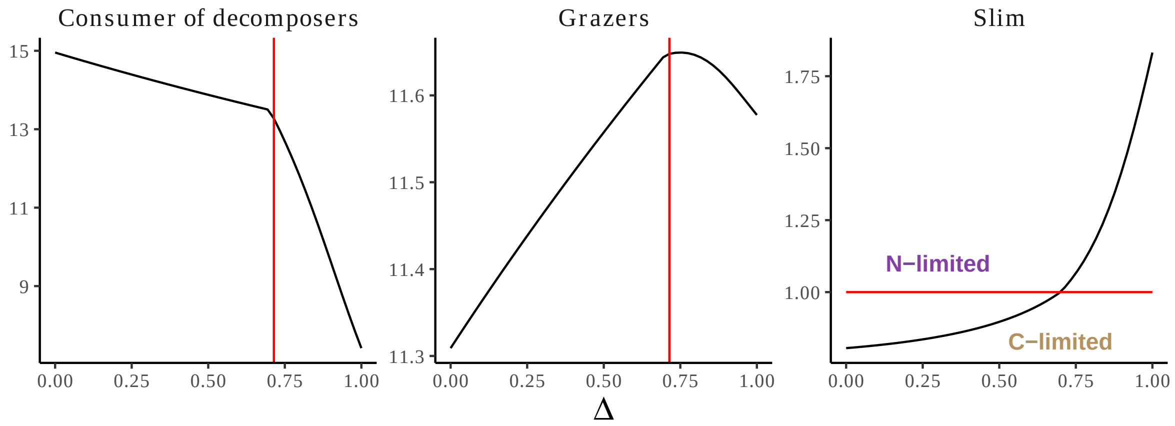


Figure S1.5: **Switch in decomposers limitation from carbon to nitrogen with increasing meta-ecosystem connectivity.**
When decomposers threshold limitation ($S_{lim}$) are near the threshold elemental ratio of 1, an increase in exchange of resources between ecosystems can relax carbon limitation until decomposers become nitrogen-limited (above red line). We also show the consumer of decomposers and grazers carbon stocks at equilibrium and how their dynamics change with the limitation of decomposers. To see this switch in decomposers limitation by spatial flows, we set $I_{N_{\mathcal{A}}} = 2.5$, $I_{D_{\mathcal{A}}} = 10$, $l_{D_{\mathcal{A}}} = 2$, $l_{D_{\mathcal{A}}} = 1$.

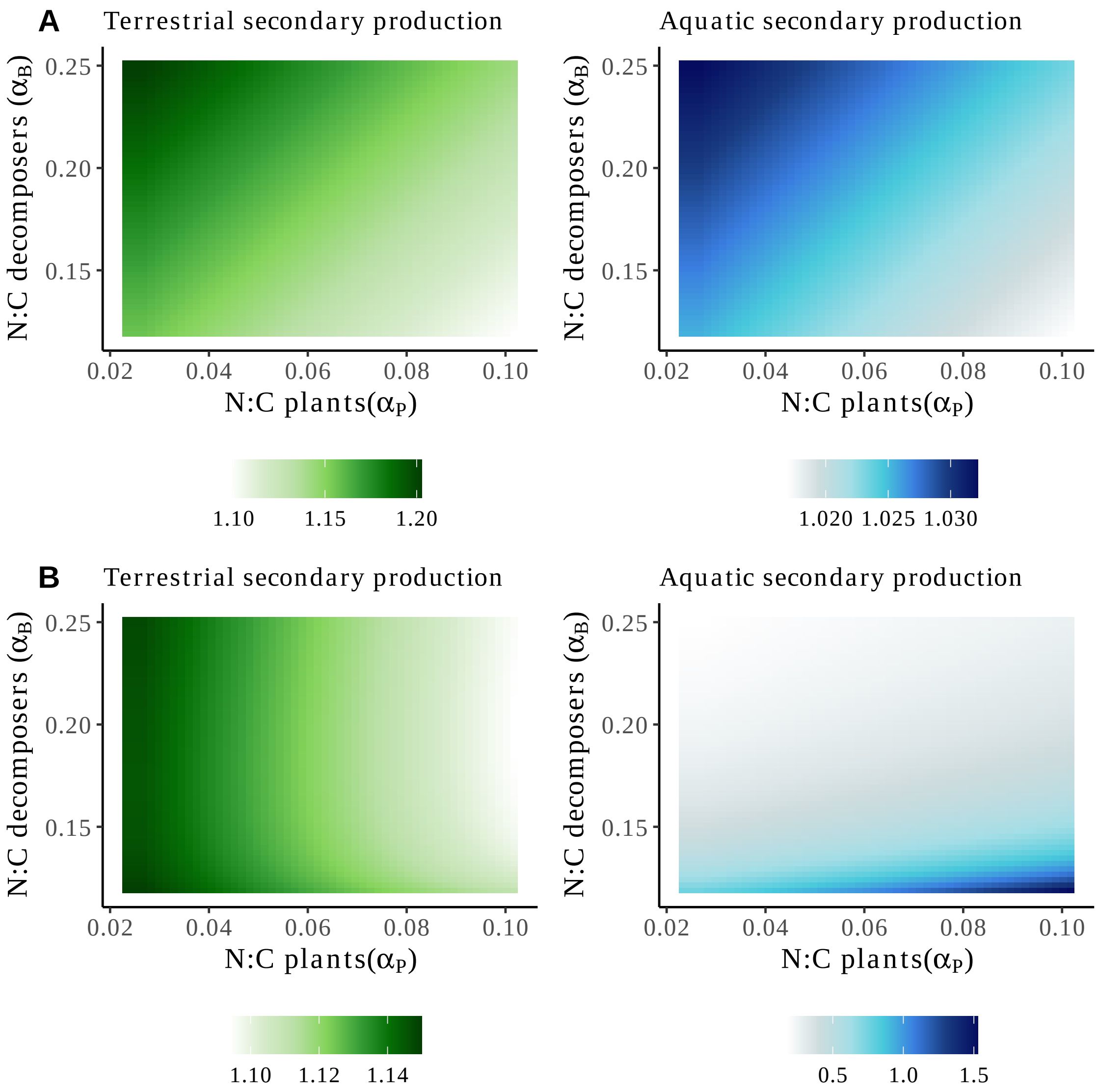


Figure S1.6: **Stoichiometry of basal species drives secondary production at terrestrial-aquatic ecotone.**

We show the variations in secondary production in aquatic (left) and terrestrial (right) ecosystems when ecosystems are connected through spatial flows ($\Delta = 1$) over the stoichiometric space of basal species (plant N:C in x-axis and decomposer N:C in y-axis). In panel (A), decomposers are carbon limited, while they are limited by nitrogen in panel (B). Production in both ecosystems is expressed in carbon units.

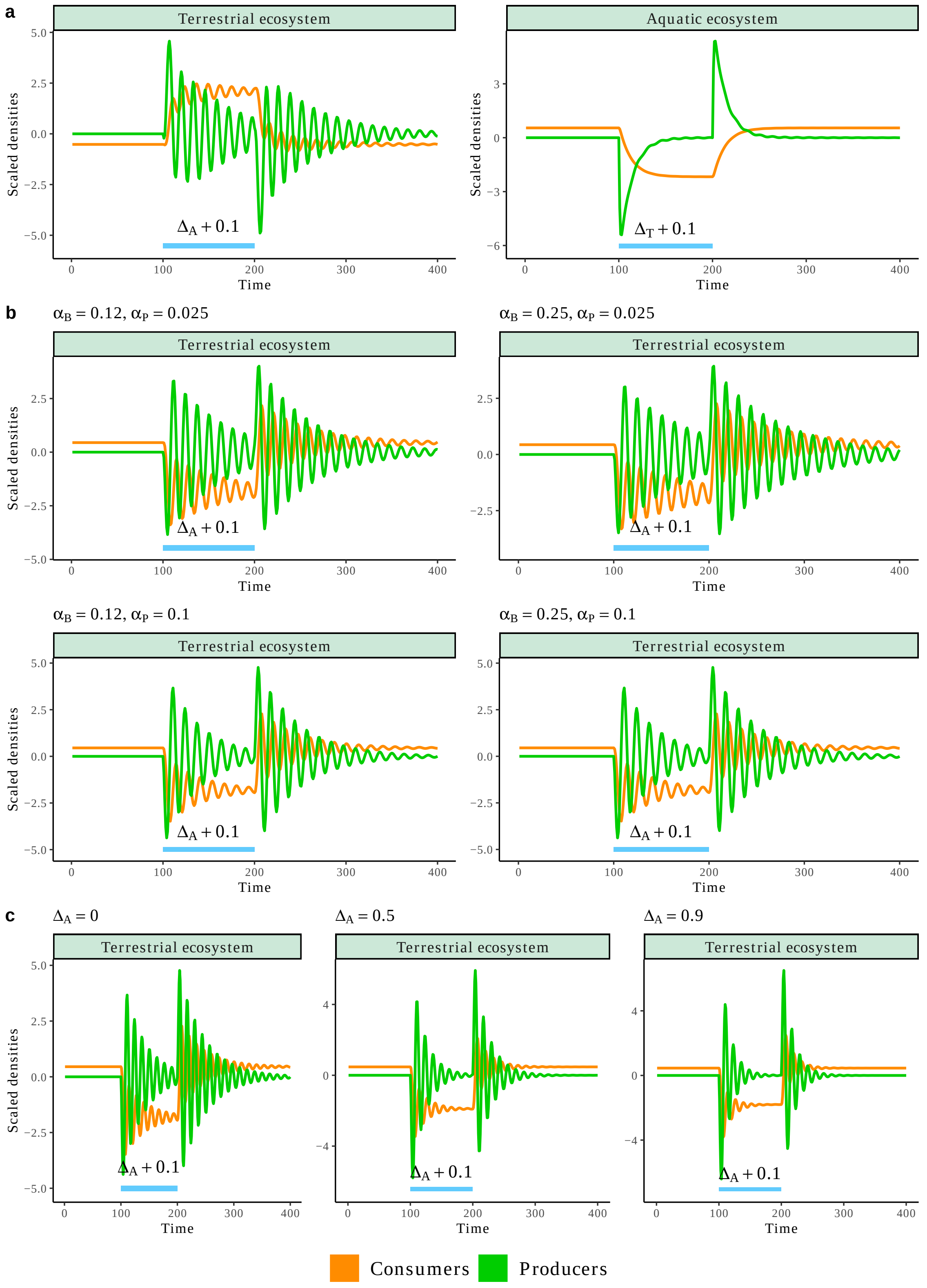


Figure S1.7: Caption on next page

**Caption for Fig. S1.7. Evaluating the spatial subsidies effect through the short-term response of ecosystems.**

Starting from meta-ecosystem equilibrium, we increased the fraction of subsidies exported to the aquatic (resp. terrestrial) ecosystem, $\Delta_T = \{\Delta_P, \Delta_G\}$ (resp. $\Delta_T = \{\Delta_B, \Delta_C\}$) by 0.1 ($\Delta_{\text{transient}} = \Delta + 0.1$) between $t = 100$ and $t = 200$ (blue rectangle), and observed the response in terrestrial (resp. aquatic) ecosystem.

(a) Response of aquatic (left) and terrestrial (right) ecosystems to a transient increase in terrestrial (left) and aquatic (right) subsidies exports under C-limitation of decomposers. In both cases, consumers ($H$ & $C$), as well as decomposers increase in density. As for the long term response, complementarity of ecosystems drives an increase in trophic level densities and therefore production in both ecosystems.

(b) Response of the aquatic ecosystem to a transient increase in exports of terrestrial subsidies under N-limitation of decomposers for different stoichiometric ratio of plants ($\alpha_P \in \{0.025, 0.1\}$) and decomposers ($\alpha_B \in \{0.12, 0.25\}$). As in Fig. 5, decomposer and consumer densities decrease in 3 out of the 4 stoichiometric conditions. This is the spatial competition between ecosystems induced by a mismatch in stoichiometry. In the one case where consumers and decomposers increase in response to more subsidies from the terrestrial, the mass-effect outbalances the stoichiometric mismatch mechanisms, therefore increasing aquatic ecosystem production.

(c) Response of the aquatic ecosystem to a transient increase in exports of terrestrial subsidies under N-limitation of decomposers for 3 levels of terrestrial ecosystem inputs ($\Delta_X \in \{0, .5, .9\}$, where $X \in \{P, B, C, H\}$). This example illustrates the interaction between stoichiometric mismatch and mass-effect mechanisms. At low coupling, stoichiometric mismatches dominate the mass-effect and therefore decomposer and consumer densities decrease during the transient increase of terrestrial inputs. However, for higher fraction of spatial subsidies transferred, the effect is reverted as seen in Fig. 5: the mass-effect mechanism outbalances the stoichiometric mismatches, therefore increasing aquatic ecosystem production.

To simplify the reading, nitrogen and detritus are not represented. Other parameters: (a): $\Delta_X = 0.5$, $\alpha_P = 0.025$, $\alpha_B = 0.12$. (b): $\Delta_X = 0$. (c): $\alpha_P = 0.1$, $\alpha_B = 0.25$.

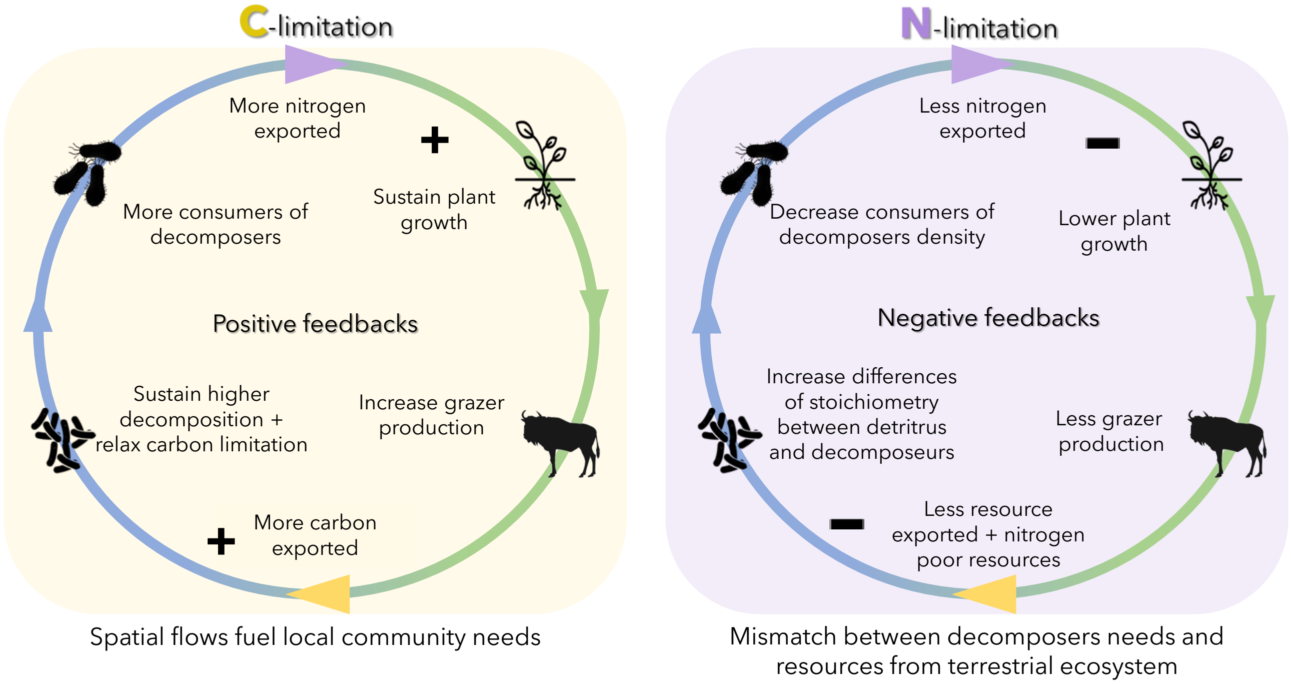


Figure S1.8: **Mechanisms and pathways of the feedbacks between aquatic and terrestrial ecosystem.** This figure does not explicitly show the mechanisms related to changes in stoichiometry of plants or decomposers.

### S2 Extracting cross-ecosystem data flows

We aimed at providing a quantitative and qualitative (N:C ratio) panorama of resource cross-ecosystem flows linking freshwater and terrestrial ecosystems. For that, we gathered in two separate databases estimates of carbon (C) and Nitrogen (N) spatial flows on the one hand and N:C ratios of resource spatial flows on the other hand. For the spatial flow database, we started from the database used in Gounand et al., 2018b providing estimates of carbon (C) flows linking many types of ecosystems, in $gC.m^{-2}.yr^{-1}$, with the area referring to the receiving ecosystem. We selected the papers reporting flows between forests, grassland, desert, agroecosystem, and stream or lake. This led to 324 values of C flow. We added the nitrogen quantification of the same flows when provided in the paper (following the same procedure as in Gounand et *al.* to convert the value in $gN.m^{-2}.yr^{-1}$), leading to 204 values for N flow. When not directly provided, we derived N flows by combining N:C ratios and C flow values. Some N:C ratios were provided in the study. If not, we used ratios of the more similar type of material possible provided by another study (*i.e.,* at best of the same species in another study; if not available of the same genus, family, order or phylum in this other). Notably, the coarser estimates (same phylum) were estimated using the C and N content in dry mass (see Table.S2.1). Finally, we completed the database with a few papers that were not in Gounand et al., 2018b but were relevant (6 papers; see references 89 to 94 in the database, containing 17 N flow values), for instance, some that only contained nutrient and not carbon flow measures.

For the stoichiometric ratio database, we collected N:C ratios of resource cross-ecosystem flows at the freshwater-terrestrial interface. We recorded the ratios present in the studies used in the spatial flow database or calculated them when both C and N flow estimates were provided in the same study (36 N:C ratios). We completed the database with twenty additional studies reporting only N:C ratios (ref 95 to 115).

In total, the two databases gather 324 C spatial flows, 204 N spatial flows, and 227 N:C ratios.

Table S2.1: **Conversion table for extracting N:C ratio from body mass.**

| Material | 1 KJ | 1g WW | 1g DW | 1g AFDW | Individual | Reference |
|---|---|---|---|---|---|---|
| Biological tissue | 0.02 gC | 0.09 gC | 0.45 gC | 0.5 gC | | Weathers et al. 2013 |
| Non woody plant detritus | | | 0.3003 gC | | | Opitz et al. 1996 |
| Terrestrial arthropods | | | 0.496 gC and 0.1 gN | | | Small et al. 2013 |
| Emergent aquatic insects (adults) | | | 0.463 gC 0.1025 gN | | | Small & Pringle 2010 |
| Emergent chironomids (adults) | | | 0.0919 gN | | | Gratton et al. 2008 |
| Amphibians (salamanders and frogs) | | | 0.44579 gC and 0.11354 gN | | | Fritz & Whiles 2018 |
| Salamanders | | | 0.4528 gC and 0.1193 gN | | | Fritz & Whiles 2018 |
| Salamander eggs and Cicadas (*Cicada magicada*) | | | 0.463 gC and 0.0892 gN 0.55086 gC and 0.10955 gN | | | Fritz & Whiles 2018 Pray et al. 2009 |
| Salmon (*Oncorhynchus nerka*) | | | | | 222.72 gC 56.072 gN | Mathiesen et al. 1988 |

Below we summarize the empirical data in tables (Table. S2.2 for stoichiometric ratio and Table. S2.3 for carbon and nitrogen flows).

Table S2.2: **Summary of stoichiometric ratio of subsidies exported at terrestrial-freshwater ecotone.**
q25, q50 and q75 being the first, second (median) and third quantiles of the N:C ratio of flows (molar) respectively. n is the number of data points. The data references as available in the Zenodo link.

| Ecosystem | n | min | q25 | q50 | q75 | max | Data references |
|---|---|---|---|---|---|---|---|
| Forest | 122 | 0.00591 | 0.0187 | 0.02674 | 0.03846 | 0.2646 | 64, 88, 91, 110, 114, 115 |
| Grassland | 46 | 0.00952 | 0.01715 | 0.15129 | 0.18317 | 0.26316 | 91, 37, 71, 24, 26, 96, 81, 97, 98, 99, 30, 57, 82, 58, 103, 104, 105, 106, 107, 111, 112, 115 |
| Lake | 15 | 0.10532 | 0.1756 | 0.18182 | 0.20137 | 0.26312 | 37, 96, 101, 102, 111, 113 |
| Stream | 35 | 0.14925 | 0.18258 | 0.20152 | 0.2146 | 0.31075 | 95, 97, 100, 74, 108, 109, 111 |

Table S2.3: **Summary of carbon and nitrogen flows at terrestrial-freshwater ecotone.** q25, q50 and q75 being the first, second (median) and third quantiles respectively. n is the number of data points. The data references as available in the Zenodo link. Fresh. = Freshwater and Terr. = Terrestrial

| Direction | Resource | n | min | q25 | q50 | q75 | max | Data references |
|---|---|---|---|---|---|---|---|---|
| Fresh. to terr. | C | 105 | 0.00224 | 0.21722 | 1.00621 | 6.00359 | 467.87459 | 1, 10, 11, 19, 23, 33, 34, 35, 36, 38, 39, 40, 41, 42, 43, 44, 45, 56, 61, 65, 66, 67, 70, 72, 73, 75, 77, 78, 79, 80, 84, 87 |
| Fresh. to terr. | N | 104 | 0.00046 | 0.04041 | 0.18643 | 0.80139 | 79.30078 | 1, 10, 11, 19, 23, 33, 34, 35, 36, 38, 39, 40, 89, 42, 43, 44, 45, 56, 61, 67, 70, 72, 73, 75, 77, 78, 117, 80, 84, 87, 91, 65 |
| Terr. to fresh. | C | 208 | 0.10532 | 3.97656 | 47.125 | 191.175 | 2085.756 | 2, 3, 4, 5, 6, 7, 8, 9, 12, 13, 14, 15, 16, 17, 18, 20, 21, 22, 24, 25, 26, 27, 28, 29, 30, 31, 32, 36, 37, 39, 45, 46, 47, 48, 49, 50, 51, 52, 53, 54, 55, 56, 57, 58, 59, 60, 62, 63, 64, 65, 68, 69, 71, 74, 76, 81, 82, 83, 85, 86, 88 |
| Terr. to fresh. | N | 93 | 0.00624 | 0.29661 | 0.992 | 2.711 | 208 | 3, 4, 9, 22, 24, 26, 30, 31, 36, 37, 39, 49, 53, 56, 57, 58, 64, 68, 69, 71, 81, 82, 83, 85, 88, 90, 91, 92, 93, 65, 94 |

### S3 Derivation of $\phi_I$ and $\phi_D$, full model system and parameters values

Table S3.1: **State variables considered in the meta-ecosystem model.**

| Symbol | Unit | Meaning |
|---|---|---|
| $B_{N_\mathcal{A}}$ | N | Nitrogen content in the decomposers |
| $B_{C_\mathcal{A}}$ | C | Carbon content in the decomposers |
| $C_{C_\mathcal{A}}$ | C | Carbon content in the consumers of decomposers |
| $P_{C_\mathcal{T}}$ | C | Carbon content in the plants |
| $G_{C_\mathcal{T}}$ | C | Carbon content in the grazers |
| $D_{N_\mathcal{A}}$ | C | Nitrogen content in the aquatic detritus |
| $D_{C_\mathcal{A}}$ | C | Carbon content in the aquatic detritus |
| $D_{N_\mathcal{T}}$ | N | Nitrogen content in the terrestrial detritus |
| $D_{C_\mathcal{T}}$ | C | Carbon content in the terrestrial detritus |
| $N_\mathcal{A}$ | N | Nitrogen stock in the aquatic ecosystem |
| $N_\mathcal{T}$ | N | Nitrogen stock in the terrestrial ecosystem |

N:C ratios of organisms were constrained using published stoichiometric data : N:C of plants and decomposers varied between 0.025-0.1 and 0.12-0.25 respectively (Elser et al., 2000; Cleveland & Liptzin, 2007; Buchkowski et al., 2019). We fixed the N:C ratio of consumers to 0.1 as no major differences between grazers and consumers of decomposer have been documented. We used two different sets of parameters in order to understand the interactions and processes at the meta-ecosystem scale under carbon and nitrogen limitation of decomposers. The two parameter sets can be found in Table S3.3. We chose to vary the fractions of subsidies that were locally or regionally recycled ($\Delta$) and the stoichiometry of organisms to account for both quantity and quality of subsidies (Sitters et al., 2015). Note that changing the stoichiometry of organisms directly changes their nitrogen content but not necessarily their carbon content.

Table S3.2: **Parameters meaning and symbol.**
The values are given for the C- and N-limited scenarios respectively

| Parameter | Unit | Meaning |
|---|---|---|
| **Terrestrial ecosystem** | | |
| $I_{N_{\mathcal{T}}}$ | N.day$^{-1}$ | Nitrogen inflow in terrestrial ecosystem |
| $I_{D_{\mathcal{T}}}$ | C.day$^{-1}$ | Detritus inflow in terrestrial ecosystem |
| $l_{D_{\mathcal{T}}}$ | day$^{-1}$ | Loss rate of detritus in terrestrial ecosystem |
| $l_{N_{\mathcal{T}}}$ | day$^{-1}$ | Loss rate of nitrogen in terrestrial ecosystem |
| $m_{\mathcal{T}}$ | day$^{-1}$ | Mineralization rate in terrestrial ecosystem |
| $e_G$ | dimensionless | Growth efficiency of grazers |
| $a_P$ | N$^{-1}$.$day^{-1}$ | Nitrogen consumption rate of plants |
| $a_G$ | C$^{-1}$.$day^{-1}$ | Consumption rate of grazers on plants |
| $d_G$ | day$^{-1}$ | Loss rate of grazers |
| $d_P$ | day$^{-1}$ | Loss rate of plants |
| **Aquatic ecosystem** | | |
| $I_{N_{\mathcal{A}}}$ | N.day$^{-1}$ | Nitrogen inflow in aquatic ecosystem |
| $I_{D_{\mathcal{A}}}$ | C.day$^{-1}$ | Detritus inflow in aquatic ecosystem |
| $l_{D_{\mathcal{A}}}$ | day$^{-1}$ | Loss rate of detritus in aquatic ecosystem |
| $l_{N_{\mathcal{A}}}$ | day$^{-1}$ | Loss rate of nitrogen in aquatic ecosystem |
| $m_{\mathcal{A}}$ | day$^{-1}$ | Mineralization rate in aquatic ecosystem |
| $e_B$ | dimensionless | Growth efficiency of decomposers |
| $e_C$ | dimensionless | Growth efficiency of consumers |
| $a_{BN}$ | day$^{-1}$ | Nitrogen consumption rate of decomposers |
| $a_{BD}$ | day$^{-1}$ | Detritus consumption rate of decomposers |
| $a_C$ | C$^{-1}$.$day^{-1}$ | Consumption rate of consumers |
| $d_B$ | day$^{-1}$ | Loss rate of decomposers |
| $d_C$ | day$^{-1}$ | Loss rate of consumers |
| **Spatial flows** | | |
| $\Delta_B, \Delta_C, \Delta_P, \Delta_G$ | dimensionless | Fraction of subsidies transferred to the other ecosystem |
| **Stoichiometric ratio** | | |
| $\alpha_B$ | N/C (molar) | Stoichiometric ratio of decomposers |
| $\alpha_C$ | N/C (molar) | Stoichiometric ratio of consumer |
| $\alpha_P$ | N/C (molar) | Stoichiometric ratio of plants |
| $\alpha_G$ | N/C (molar) | Stoichiometric ratio of grazers |

We performed a sensitivity analysis to make sure that our results were robust to variations in parameter values. We established the range of variation of each parameter so that *(i)* we stayed in the same scenario of decomposer limitation and *(ii)* all trophic levels were coexisting along the ranges of parameter values explored (see Table S7.1). We evaluated the quantitative effect of parameter variation by taking the range of variation of production (*i.e.,* $|max(\text{Production}) - min(\text{Production})|$) over the range of parameter values explored for two different stoichiometries of decomposers ($\alpha_B \in \{0.12, 0.25\}$) and two stoichiometries of plants ($\alpha_P \in \{0.025, 0.1\}$).

#### Derivation of $\phi_I$ and $\phi_D$

Stoichiometric homeostasis of decomposers implies :

$$\frac{dB_{N_\mathcal{A}}}{dt} = \alpha_B \frac{dB_{C_\mathcal{A}}}{dt}$$

which constrains the immobilization and decomposition flows :

$$\phi_I = \frac{\alpha_B - \alpha_D}{\alpha_B}\,\phi_D \text{ , with } \alpha_D = \frac{D_{N_\mathcal{A}}}{D_{C_\mathcal{A}}}$$

We explored in the main text two scenarios: decomposers are limited by nitrogen or by carbon. When decomposers are N-limited, decomposers immobilize nitrogen and modulate their decomposition to maintain their homeostasis. Therefore: $\phi_I$ constrains $\phi_D$. To the contrary, in the C-limited case, the decomposition process ($\phi_D$) constrains the uptake or release of nitrogen ($\phi_I$). Thus, we get the following formulas for the immobilization and decomposition flows for carbon and nitrogen-limited decomposers :

| | $\phi_I$ | $\phi_D$ |
|---|---|---|
| **C-limited** | $\frac{\alpha_B - \alpha_D}{\alpha_B}\phi_D$ | $a_B D_{C_\mathcal{A}}$ |
| **N-limited** | $a_{BN} N_\mathcal{A}$ | $\frac{\alpha_B}{\alpha_B - \alpha_D}\phi_I$ |

In the nitrogen-limited environment, decomposers feed on both nitrogen and detritus. Under such limitation, a decrease in N:C ratio of detritus limits the decomposition rate of decomposers (*i.e.*, $\phi_D$ decreases due to an increase of $\alpha_B - \alpha_D$). On the contrary, when decomposers are C-limited they decompose detritus into nitrogen and immobilize (when $\alpha_B - \alpha_D > 0$) or excrete (when $\alpha_B - \alpha_D < 0$) nitrogen. At equilibrium, decomposers are C-limited (resp. N-limited) if the limitation threshold $S_{lim}$ is below (resp. above) the threshold elemental ratio (TER, Frost et al., 2006) that equals 1 :

$$\text{where } S_{\text{lim}} = \frac{(\alpha_B - \bar{\alpha}_D) a_B \bar{D}_{C_{\mathcal{A}}}}{\alpha_B a_{BN} \bar{N}_{\mathcal{A}}}, \quad \bar{\alpha}_D = \frac{\bar{D}_{N_{\mathcal{A}}}}{\bar{D}_{C_{\mathcal{A}}}}$$

, where the bar is used for equilibrium quantities.

The carbon or nitrogen limitations of decomposers might change with the stoichiometry of both decomposers and detritus and with the resource stocks at equilibrium. We assumed donor-control flows for decomposers (Cherif & Loreau, 2013; Daufresne & Loreau, 2001), meaning that decomposition and immobilization flow only depend on the quantity of resources but not on the decomposer density. Note that using Lotka-Voltera functional response for decomposers gave qualitatively the same results, as decomposers' density was constant due to top-down control by their consumers, but the species and stock dynamics were oscillating under N-limitation. We used the Liebig law, which assumes that the limiting resource is the scarcer one in the ecosystem, to express the decomposition and immobilization flows :

$$\phi_I = min(\overbrace{\frac{(\alpha_B - \alpha_D)}{\alpha_B} e_B a_{BD} D_{C_{\mathcal{A}}}}^{\text{C-limited}}, \overbrace{a_{BN} N_{\mathcal{A}}}^{\text{N-limited}})$$

$$\phi_D = min(\overbrace{e_B a_{BD} D_{C_{\mathcal{A}}}}^{\text{C-limited}}, \overbrace{(\frac{\alpha_B - \alpha_D}{\alpha_B})^{-1} a_{BN} N_A}^{\text{N-limited}})$$

#### Full model system

With the spatial flow of subsidies, the nitrogen dynamics of plants, consumers, and grazers, and both the decomposition and immobilization flows, we get the following system. First, for terrestrial ecosystem :

$$\frac{dG_{C_\mathcal{T}}}{dt} = \overbrace{e_G\ a_G P_{C_\mathcal{T}} G_{C_\mathcal{T}}}^{\text{Consumption of grazers (carbon)}} - \overbrace{d_G G_{C_\mathcal{T}}}^{\text{Loss of carbon by grazers}}$$

$$\frac{dG_N}{dt} = \overbrace{e_G\ a_G P_{C_\mathcal{T}} G_{C_\mathcal{T}} \alpha_G - d_G G_{C_\mathcal{T}} \alpha_G}^{\text{Same but for nitrogen part}}$$

$$\frac{dP_{C_\mathcal{T}}}{dt} = \overbrace{a_P N_\mathcal{T} P_{C_\mathcal{T}}}^{\text{Photosynthesis}} - \overbrace{a_G P_{C_\mathcal{T}} G_{C_\mathcal{T}}}^{\text{Consumption by grazers}} - \overbrace{d_P P_{C_\mathcal{T}}}^{\text{Loss of carbon by plants}}$$

$$\frac{dP_N}{dt} = \overbrace{a_P N_\mathcal{T} P_{C_\mathcal{T}} \alpha_P - a_G P_{C_\mathcal{T}} G_{C_\mathcal{T}} \alpha_P - d_P P \alpha_P}^{\text{Same but for nitrogen part}}$$

$$\begin{aligned}\frac{dD_{N_\mathcal{T}}}{dt} = {} & \overbrace{I_{D_\mathcal{T}} \frac{D_{N_\mathcal{T}}}{D_{C_\mathcal{T}}}}^{\text{Inflow of detritus}} - \overbrace{l_{D_\mathcal{T}} D_{N_\mathcal{T}}}^{\text{Leaching of detritus}} + \overbrace{\alpha_G d_G G_{C_\mathcal{T}} (1-\Delta_G)}^{\text{Grazers-produced detritus locally recycled}} + \\ & \overbrace{\alpha_P d_P P_{C_\mathcal{T}} (1-\Delta_P)}^{\text{Plants-produced detritus locally recycled}} + \overbrace{\alpha_C d_C C_{C_\mathcal{A}} \Delta_C}^{\text{Consumers-produced detritus regionally transferred}} + \\ & \overbrace{\alpha_B d_B B_{C_\mathcal{A}} \Delta_B}^{\text{Decomposers-produced detritus regionally transferred}} - \overbrace{m_\mathcal{T} D_{N_\mathcal{T}}}^{\text{Mineralization of detritus}}\end{aligned}$$

$$\frac{dD_{C_\mathcal{T}}}{dt} = \overbrace{I_{D_\mathcal{T}} - l_{D_\mathcal{T}} D_{C_\mathcal{T}} + d_G G_{C_\mathcal{T}} (1-\Delta_G) + d_P P_{C_\mathcal{T}} (1-\Delta_P) + d_C C_{C_\mathcal{A}} \Delta_C + d_B B_{C_\mathcal{A}} \Delta_B - m_\mathcal{T} D_{C_\mathcal{T}}}^{\text{Same as above but for the carbon part of detritus}}$$

$$\begin{aligned}\frac{dN_\mathcal{T}}{dt} = {} & \overbrace{I_{N_\mathcal{T}}}^{\text{Nitrogen inflow}} - \overbrace{l_{N_\mathcal{T}} N_\mathcal{T}}^{\text{Leaching of nitrogen}} + \overbrace{(\alpha_P - e_G \alpha_G) a_G P_{C_\mathcal{T}} G_{C_\mathcal{T}} (1-\Delta_G)}^{\text{Nitrogen from stoichiometric imbalance of grazers}} + \\ & \overbrace{(\alpha_B - e_C \alpha_C) a_C B_{C_\mathcal{A}} C_{C_\mathcal{A}} \Delta_C}^{\text{Nitrogen from stoichiometric imbalance of consumers of decomposers}} \\ & - \overbrace{\alpha_P a_P N_\mathcal{T} P_{C_\mathcal{T}}}^{\text{Nitrogen uptake from plants}} + \overbrace{m_\mathcal{T} D_{N_\mathcal{T}}}^{\text{Mineralization of detritus into nitrogen}}\end{aligned}$$

And for aquatic ecosystem :

$$\frac{dC_{C_\mathcal{A}}}{dt} = \overbrace{e_C\ a_C B_{C_\mathcal{A}} C_{C_\mathcal{A}}}^{\text{Consumption by consumers (carbon)}} - \overbrace{d_C C_{C_\mathcal{A}}}^{\text{Loss of carbon by consumers}}$$

$$\frac{dC_N}{dt} = \overbrace{e_C\ a_C B_{C_\mathcal{A}} C_{C_\mathcal{A}} \alpha_C - d_C C_{C_\mathcal{A}} \alpha_C}^{\text{Same but for nitrogen part}}$$

$$\frac{dB_{C_\mathcal{A}}}{dt} = \overbrace{\phi_D}^{\text{Decomposition of detritus (carbon)}} - \overbrace{d_B B_{C_\mathcal{A}}}^{\text{Loss of carbon by decomposers}} - \overbrace{a_C B_{C_\mathcal{A}} C_{C_\mathcal{A}}}^{\text{Consumption by consumers (carbon)}} - \overbrace{m_\mathcal{A} B_{C_\mathcal{A}}}^{\text{Heterotrophic respiration}}$$

$$\frac{dB_{N_\mathcal{A}}}{dt} = \overbrace{\phi_I \alpha_B}^{\text{Immobilization of detritus}} + \overbrace{\phi_D \frac{D_{N_\mathcal{A}}}{D_{C_\mathcal{A}}}}^{\text{Decomposition of detritus (nitrogen)}} - \overbrace{a_C B_{C_\mathcal{A}} C_{C_\mathcal{A}} \alpha_B}^{\text{Consumption by consumers (nitrogen)}} - \overbrace{d_B B_{C_\mathcal{A}} \alpha_B}^{\text{Loss of nitrogen by decomposers}} - \overbrace{m_\mathcal{A} B_{C_\mathcal{A}} \alpha_B}^{\text{Nutrient mineralization}}$$

$$\frac{dD_{N_\mathcal{A}}}{dt} = \overbrace{I_{D_\mathcal{A}} \frac{D_{N_\mathcal{A}}}{D_{C_\mathcal{A}}}}^{\text{Inflow of detritus}} - \overbrace{l_{D_\mathcal{A}} D_{N_\mathcal{A}}}^{\text{Leaching of detritus}} + \overbrace{\alpha_C d_C C_{C_\mathcal{A}}(1-\Delta_C)}^{\text{Consumers-produced detritus locally transferred}} + \overbrace{\alpha_B d_B B_{C_\mathcal{A}}(1-\Delta_B)}^{\text{Decomposers-produced detritus locally transferred}} + \overbrace{\alpha_G d_G G_{C_\mathcal{T}} \Delta_G}^{\text{Grazers-produced detritus regionally recycled}} + \overbrace{\alpha_P d_P P_{C_\mathcal{T}} \Delta_P}^{\text{Plants-produced detritus regionally recycled}} - \overbrace{\phi_D \frac{D_{N_\mathcal{A}}}{D_{C_\mathcal{A}}}}^{\text{Decomposition of detritus}}$$

$$\frac{dD_{C_\mathcal{A}}}{dt} = \overbrace{I_{D_\mathcal{A}} - l_{D_\mathcal{A}} D_{C_\mathcal{A}} + d_C C_{C_\mathcal{A}}(1-\Delta_C) + d_B B_{C_\mathcal{A}}(1-\Delta_B) + d_G G_{C_\mathcal{T}} \Delta_G + d_P P_{C_\mathcal{T}} \Delta_P - \phi_D}^{\text{Same but for carbon}}$$

$$\frac{dN_\mathcal{A}}{dt} = \overbrace{I_{N_\mathcal{A}}}^{\text{Nitrogen inflow}} - \overbrace{l_{N_\mathcal{A}} N_\mathcal{A}}^{\text{Leaching of nitrogen}} + \overbrace{(\alpha_B - e_C\alpha_C) a_C B_{C_\mathcal{A}} C_{C_\mathcal{A}} (1-\Delta_C)}^{\text{Nitrogen from stoichiometric imbalance of consumers of decomposers}} + \overbrace{(\alpha_P - e_G\alpha_G) a_G P_{C_\mathcal{T}} G_{C_\mathcal{T}} \Delta_G}^{\text{Nitrogen from stoichiometric imbalance of grazers}} - \overbrace{\phi_I \alpha_B}^{\text{Immobilization by decomposers}} + \overbrace{m_\mathcal{A} B_{C_\mathcal{A}} \alpha_B}^{\text{Mineralization of detritus into nitrogen by decomposers}}$$

$$\phi_I = min((\frac{\alpha_B - \frac{D_{N_\mathcal{A}}}{D_{C_\mathcal{A}}}}{\alpha_B}) e_B a_{BD} D_{C_\mathcal{A}}, a_{BN} N_\mathcal{A})$$

$$\phi_D = min(e_B a_{BD} D_{C_\mathcal{A}}, (\frac{\alpha_B - \frac{D_{N_\mathcal{A}}}{D_{C_\mathcal{A}}}}{\alpha_B})^{-1} a_{BN} N_A)$$

#### Details on the simulation method

The model was run over a long time (10000-time steps), which was more than enough to converge to the equilibrium of each trophic level and resources (with Type I functional or donor-controlled responses, the systems typically converge in about 1000 to 2000 time steps). Running the model for such a long time is necessary to compute the feedbacks. A trophic level (*resp.*, a resource) was considered extinct (*resp.* empty) if its value was below $10^{-5}$.

#### Parameter values

We focused our analysis on the qualitative behavior that could emerge in ecosystems coupled by spatial flows with different stoichiometric compositions, rather than exploring the full range of parameter values. By compiling data from the literature, we found values of N:C to be lower in aquatic consumers than in their resource (bacterial decomposers), while N:C is typically higher in grazers than in plants (Fig. 1B). Nevertheless, we performed analyses on the parameter values (Appendix S7), the structure of the trophic chains (Appendix S4), and the functional responses (Appendix S6). Functional response parameters were taken from the literature and some of them were assumed to allow coexistence (all resource stocks and trophic levels having a positive value) in each isolated ecosystem. Note that coexistence was facilitated by the inflow of detritus in the aquatic ecosystem ($I_{D_{\mathcal{A}}}$) that generates a continuous input of carbon in the aquatic ecosystem. More precisely, parameters captured the differences in energy transfer and primary productivity between net heterotrophic and net autotrophic ecosystems (*i.e.* forest doing more primary production while streams are more efficient to transfer energy up to the higher trophic levels; Shurin et al., 2006; Gounand et al., 2020; Harvey et al., 2021). Therefore, we set $a_C > a_G$ and $e_C > e_G$. Similarly, we accounted for differences in mineralization rates, with aquatic ecosystems being more efficient to mineralize organic matter compared to terrestrial ones ($m_{\mathcal{T}} < m_{\mathcal{A}}$;Gounand et al., 2020). There are two parameter sets for C- and N-limitation. These parameter sets were also chosen so that decomposers remained in the same limitation all over the range of basal species stoichiometry.

Table S3.3: **Parameters values for the simulations performed.**
The values are given for the C- and N-limited scenarios respectively

| Class | Parameter | Value | Source |
|---|---|---|---|
| **Terrestrial ecosystem** | $I_{N_{\mathcal{T}}}$ | 7 | Assumed value |
| | $I_{D_{\mathcal{T}}}$ | 7 | Assumed value |
| | $l_{D_{\mathcal{T}}}$ | 1 | Assumed value |
| | $l_{N_{\mathcal{T}}}$ | 1 | Assumed value |
| | $m_{\mathcal{T}}$ | 0.1 | Assumed value |
| | $e_G$ | 0.25 | Assumed value |
| | $a_P$ | 0.34 | Cherif & Loreau, 2013 |
| | $a_G$ | 0.2 | Assumed value |
| | $d_G$ | 0.1 | Attayde & Ripa, 2008 |
| | $d_P$ | 0.1 | Attayde & Ripa, 2008 |
| **Aquatic ecosystem** | $I_{N_{\mathcal{A}}}$ | 7, 2 | Assumed values |
| | $I_{D_{\mathcal{A}}}$ | 7, 12 | Assumed values |
| | $l_{D_{\mathcal{A}}}$ | 1 | Assumed value |
| | $l_{N_{\mathcal{A}}}$ | 1 | Assumed value |
| | $m_{\mathcal{A}}$ | 0.5 | Zelnik et al., 2021 |
| | $e_B$ | 0.5 | del Giorgio & Cole, 1998 |
| | $e_C$ | 0.5 | Zelnik et al., 2021 |
| | $a_{BN}$ | 1, 0.25 | Zou et al., 2016 |
| | $a_{BD}$ | 0.83 | Boit et al., 2012 |
| | $a_C$ | 0.3 | Assumed value |
| | $d_B$ | 0.1 | Attayde & Ripa, 2008 |
| | $d_C$ | 0.1 | Assumed value |
| **Spatial flows** | $\Delta_B, \Delta_C, \Delta_P, \Delta_G$ | [0,1] | vary |
| **Stoichiometric ratio** | $\alpha_B$ | [0.12-0.25] | Cleveland & Liptzin, 2007 |
| | | | Buchkowski et al., 2019 |
| | $\alpha_C$ | | Elser et al., 2000 |
| | | | Martinson et al., 2008 |
| | $\alpha_P$ | | Cleveland & Liptzin, 2007 |
| | | | Buchkowski et al., 2019 |
| | | | Mcgroddy et al., 2004 |
| | $\alpha_G$ | | Elser et al., 2000 |
| | | | Martinson et al., 2008 |

### S4 Adding a trophic level

In this section, we relaxed the hypothesis on the structure of the two ecosystems by adding a top predator in both the net autotrophic ($T_G$) and net heterotrophic ecosystems ($T_C$). Top predators of grazers and consumers of decomposers ($T_G$ and $T_C$ respectively) consume primary consumers at a rate $a_{TG}$ and $a_{TC}$ but only a fraction $e_{TG}, e_{TC}$ of the ingested food is assimilated. We considered that the stoichiometry of these top predators is similar between ecosystems ($\alpha_{TG} = \alpha_{TC} = 0.1$). Both top predators are held at a fixed stoichiometry. As primary consumers, top predators excrete the excess of nitrogen due to stoichiometric imbalance between their resource need and their prey stoichiometry: rate $\beta_{Ti} f_{Ti}(i, Ti)$, where $\beta_{Ti} = (\alpha_i - e_{Ti}\alpha_{Ti})$ and $i \in \{G, C\}$. Finally, each top predator has its decay rate $d_{TG}$ or $d_{TC}$. The detritus produced also fuels the detritus pool of each ecosystem. The results are presented in Figs. S4.1-4, and show qualitatively similar results compared to the ones in the main text. Interestingly, in a C-limited scenario when plants and decomposers are no longer controlled by their respective consumers, plants reach high biomass and seem to drive the effects at the meta-ecosystem scale (*i.e.,* not much variation is observed along the stoichiometric ratio of decomposers). In the N-limited scenario, the stoichiometric ratio of decomposers has a drastic impact in exacerbating (low $\alpha_B$) or reducing (high $\alpha_B$) the stoichiometric mismatch with the detritus. This drives the patterns observed in Fig. S4.2-left: productions in the aquatic ecosystem is maximized when the stoichiometric mismatch between decomposers and their detritus is low (low $\alpha_B$), and when terrestrial ecosystem export carbon poor plant subsidies (low $\alpha_P$).

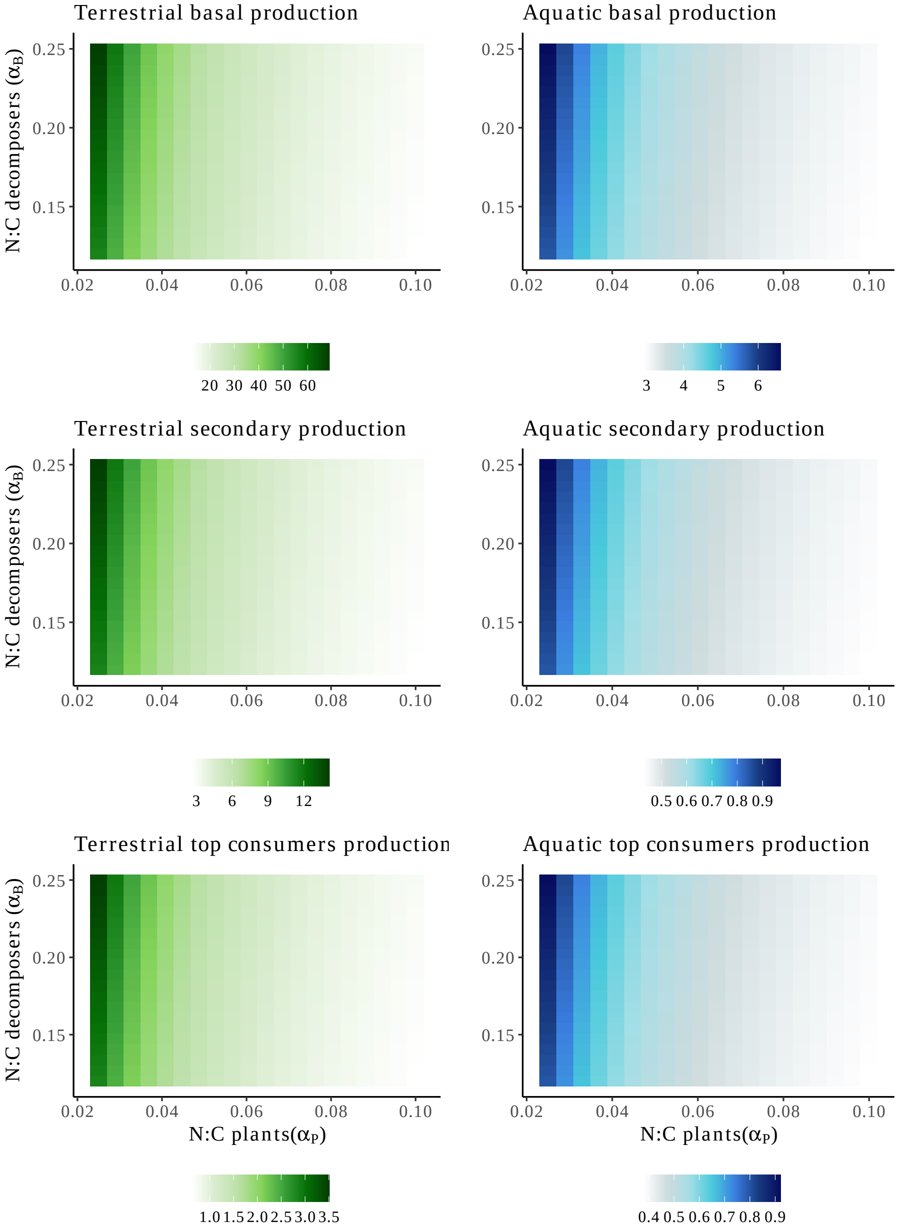


Figure S4.1: **Sensitivity analysis of the food-web structure: C-limited decomposers.**. We performed sensitivity analysis on the food-webs structure by adding top predators in both ecosystems. We measured the primary, secondary, and top production in both terrestrial and aquatic ecosystems. The parameters used are the same as in the C-limited scenario with an exception for $a_P$, $a_G$, and $a_C$ which were set to 0.1, 0.4, and 0.5 respectively to allow coexistence. Top predator parameters were chosen so that we keep coexistence for the range of stoichiometric parameters explored: $a_{TC}$=0.1, $e_{TC}$=1, $a_{TG}$=0.5, $e_{TG}$=0.25, $d_{TC} = 0.05$ and $d_{TH} = 0.15$. Here decomposers are carbon limited.

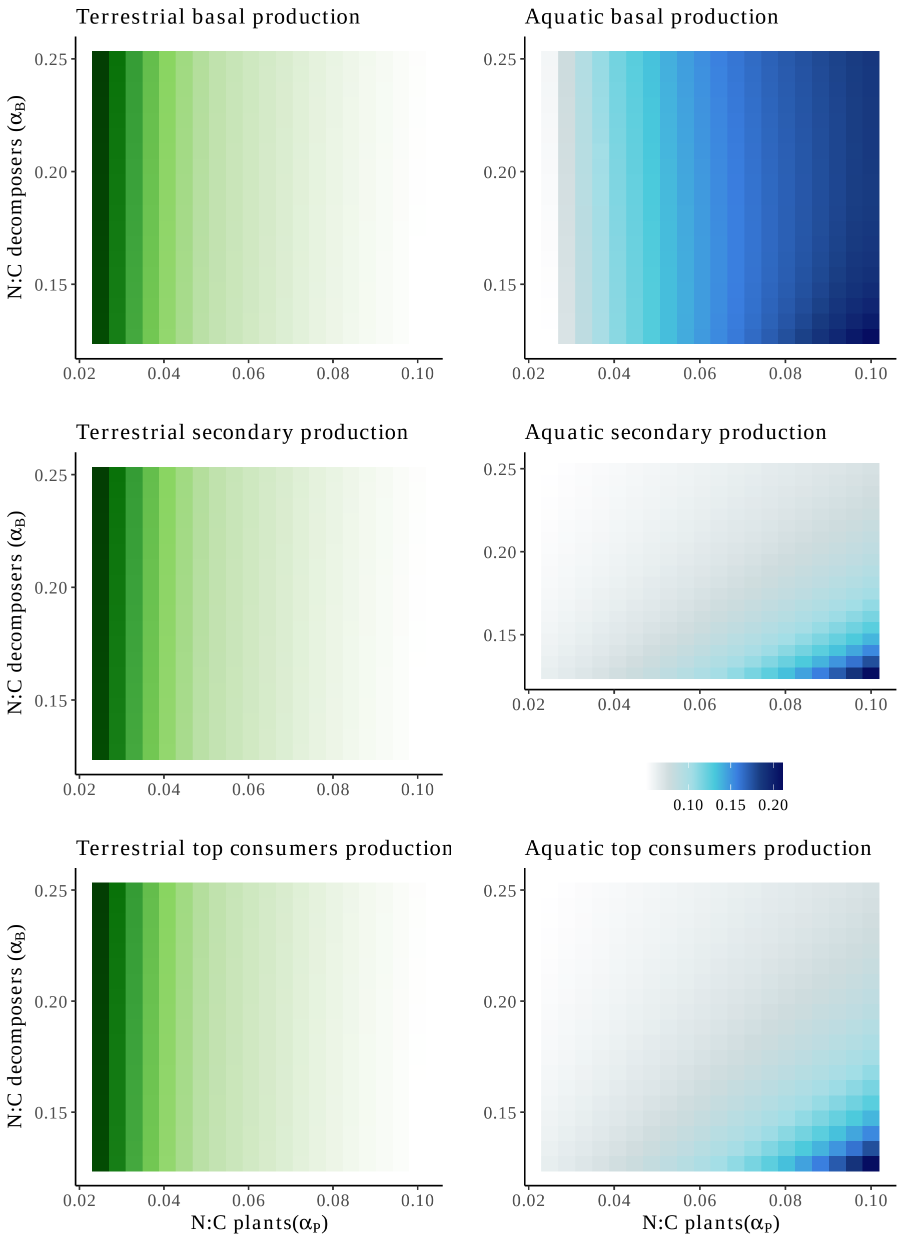


Figure S4.2: **Sensitivity analysis of the food-web structure: N-limited decomposers..** The legend is the same as in Fig. S4.1. We set: $I_{N_{\mathcal{A}}}$=5, $l_{N_{\mathcal{G}}}$=2, $I_{D_{\mathcal{A}}}$=12, $l_{D_{\mathcal{A}}}$=1.5 and $a_{BN}$=0.1 such that decomposers are nitrogen limited.

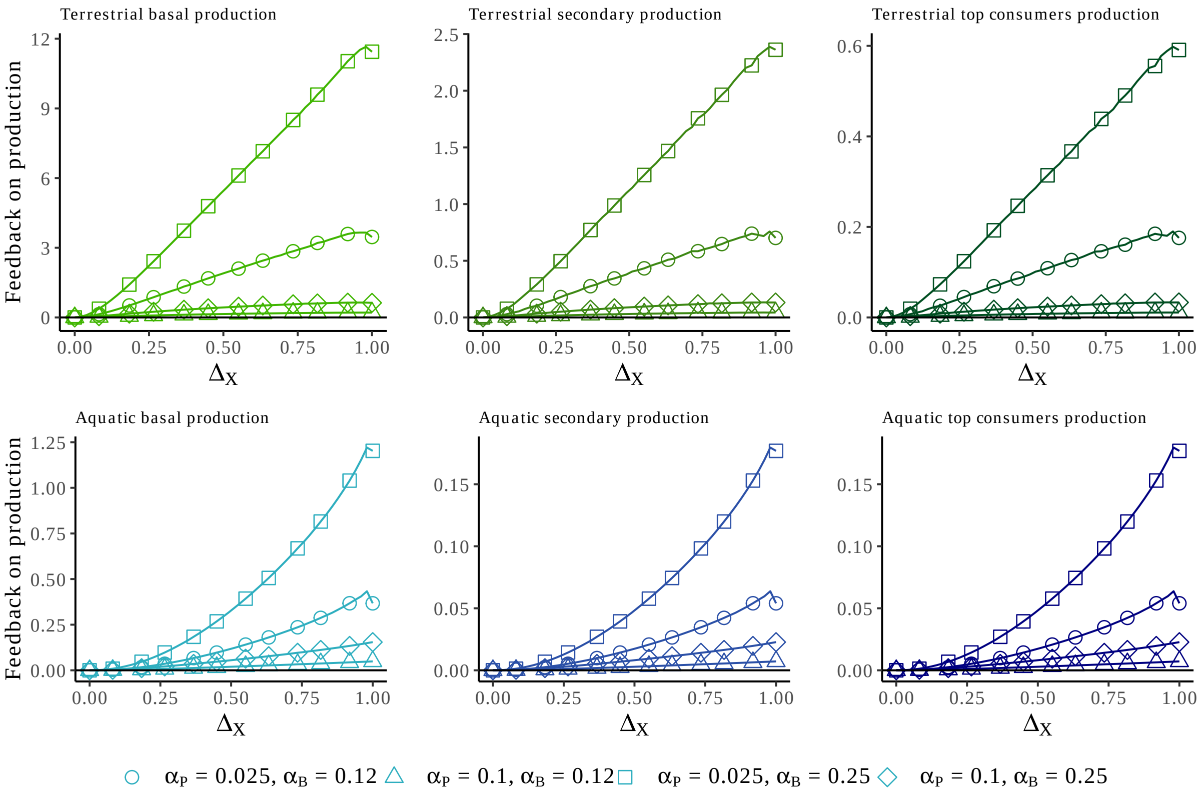


Figure S4.3: **Sensitivity analysis of the food-web structure: feedbacks under C-limited decomposers.**.

We computed the feedback as shown in Fig. S1.1 and Eq. 4. The feedback is computed for each trophic level (columns) in both ecosystems (rows). Here decomposers are limited by carbon.

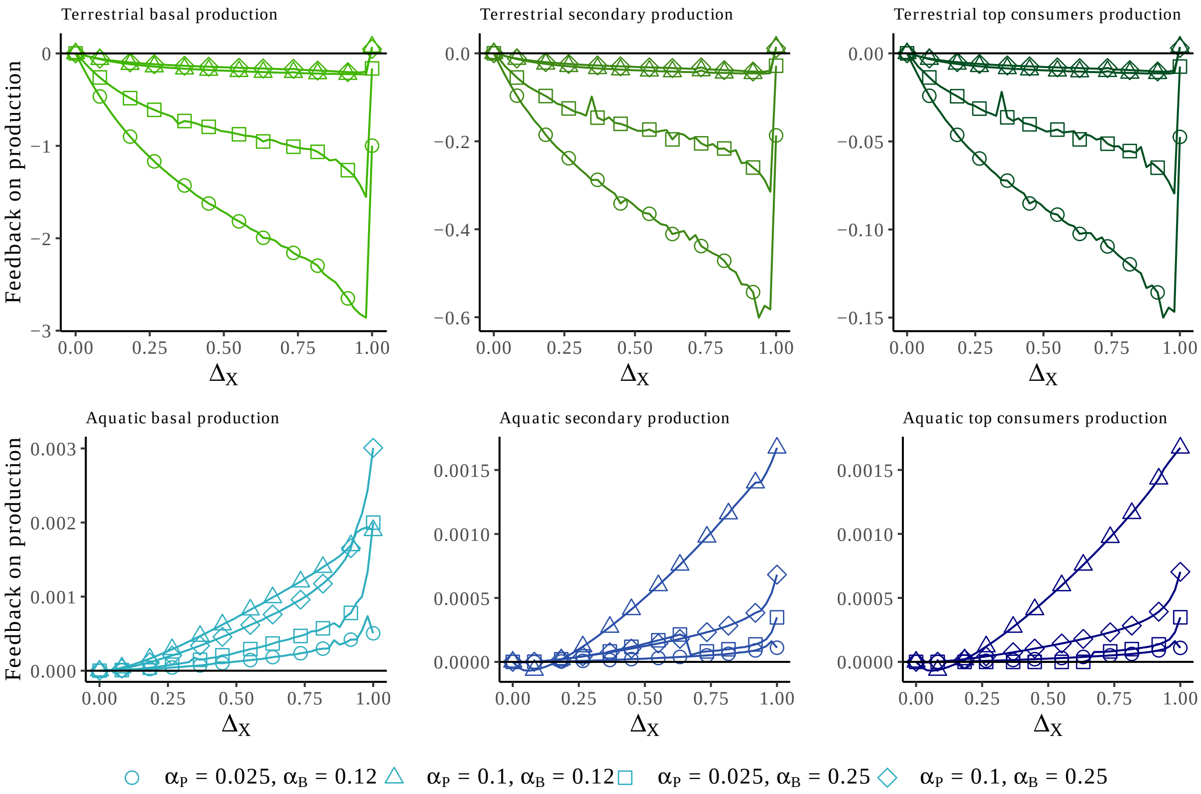


Figure S4.4: **Sensitivity analysis of the food-web structure: feedbacks under N-limited decomposers.**.

We computed the feedback as shown in Fig. S1.1 and Eq. 4. The feedback is computed for each trophic level (columns) in both ecosystems (rows). Here decomposers are limited by nitrogen.

### S5 Co-limitation of decomposers

In the main text, we restricted the analysis to the case where a strict limitation of carbon or nitrogen was observed in decomposer populations. Here we relax this hypothesis by assuming that decomposers are co-limited by nitrogen and carbon. In fact, co-limitation is expected to be selected at the community scale due to competitive exclusion that favors the most competitive species for nitrogen (plants or decomposers). Co-limitation has been experimentally observed (Danger et al., 2008; Daufresne et al., 2008) and previously considered in a few non-spatial theoretical models (Cherif & Loreau, 2007; Halvorson et al., 2017). Here we investigate the impact of co-limitation in a meta-ecosystem context. We considered independent co-limitation which assumes a synergy between two limiting resources (Harpole et al., 2011; Sperfeld et al., 2016). When decomposers are co-limited, the decomposition flux is defined as $\phi_D = e_B a_{BD} D_{C_A} \, a_{BN} N_A B_{C_A}$ and immobilization flux as $\phi_I = \dfrac{\alpha_B - \alpha_D}{\alpha_B} \phi_D$ so that decomposers are held at a constant stoichiometry (Sperfeld et al., 2012; Wirtz & Kerimoglu, 2016). Under the co-limitation scenario for decomposers, we chose to represent the primary production with $\phi_D$.

The parameters were the same as in the C-limited scenario (see Table S3.3).

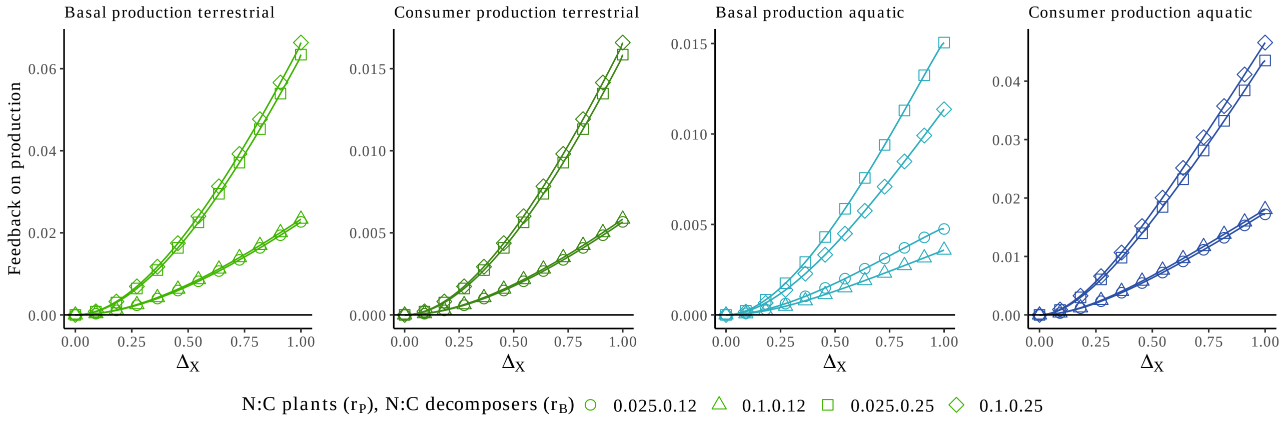


Figure S5.1: **Feedback at the landscape extent under co-limitation of decomposers.** We computed the feedback as shown in Fig. S1.1 and Eq. 4. The feedback was computed for each of the trophic levels in both ecosystems (columns) under the co-limitation of decomposers.

### S6 Donor-Control functional responses

In this section, we performed a sensitivity analysis on the functional responses used in the main text by using donor-control functional responses for each trophic interaction. In this case, the flow only depends on the size of the donor pool (*e.g.*, $f_P = a_P N_{\mathcal{T}}$ for plants). Parameters are the same as in Table S3.3. We set $l_{N_{\mathcal{A}}} = 2$ in the N-limited scenario so that decomposers stayed in nitrogen limitation for the range of stoichiometric parameters explored. The results with donor-control functional responses are qualitatively similar to the ones in the main text (see Figs. S6.1, S6.2, S6.3).

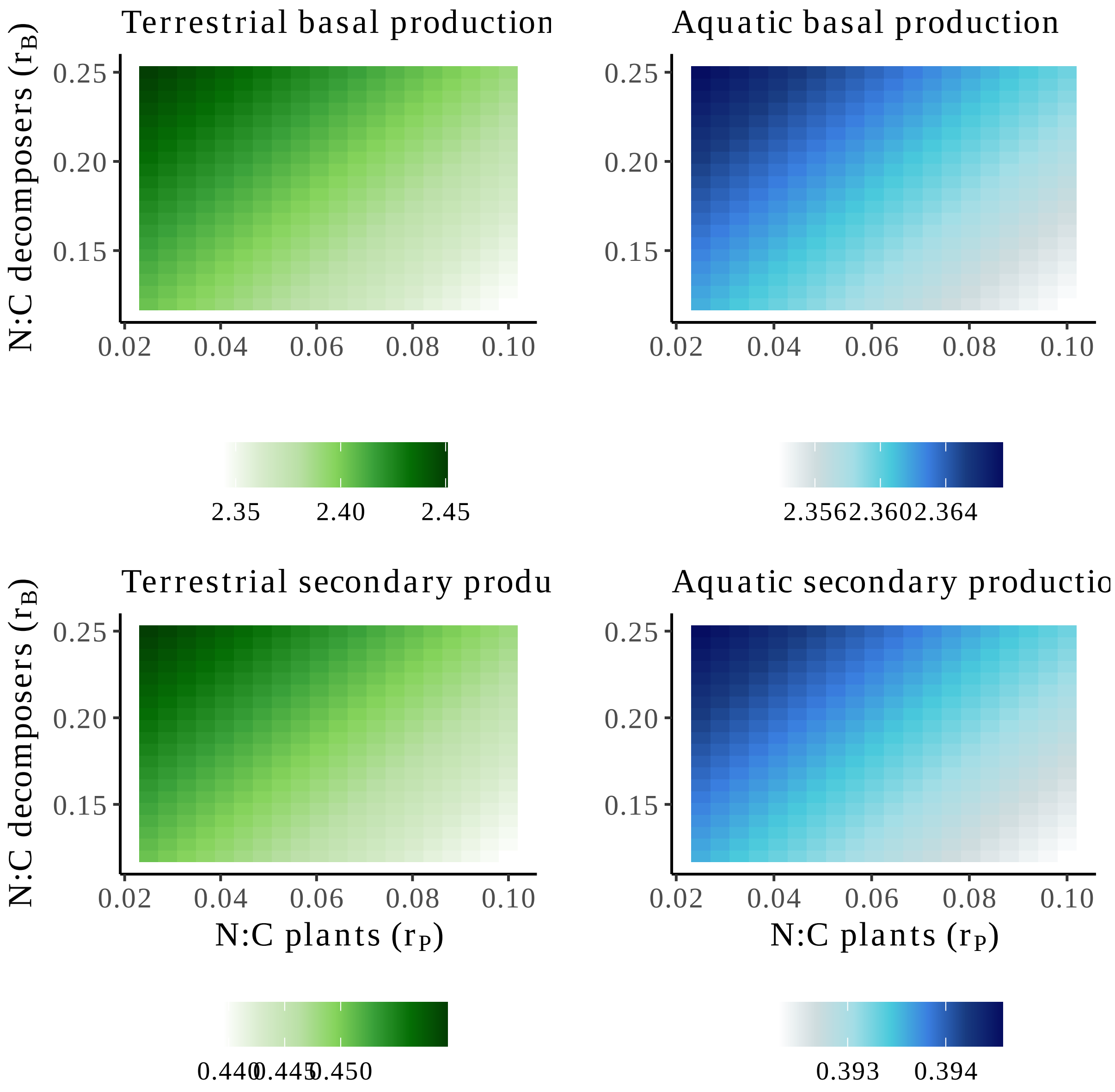


Figure S6.1: **Main conclusions are robust to donor-control functional responses under the carbon limitation of decomposers.**
The legend is the same as in Fig. 3 but we also show the secondary production of both terrestrial and aquatic ecosystems (bottom figures). Here, decomposers are carbon limited.

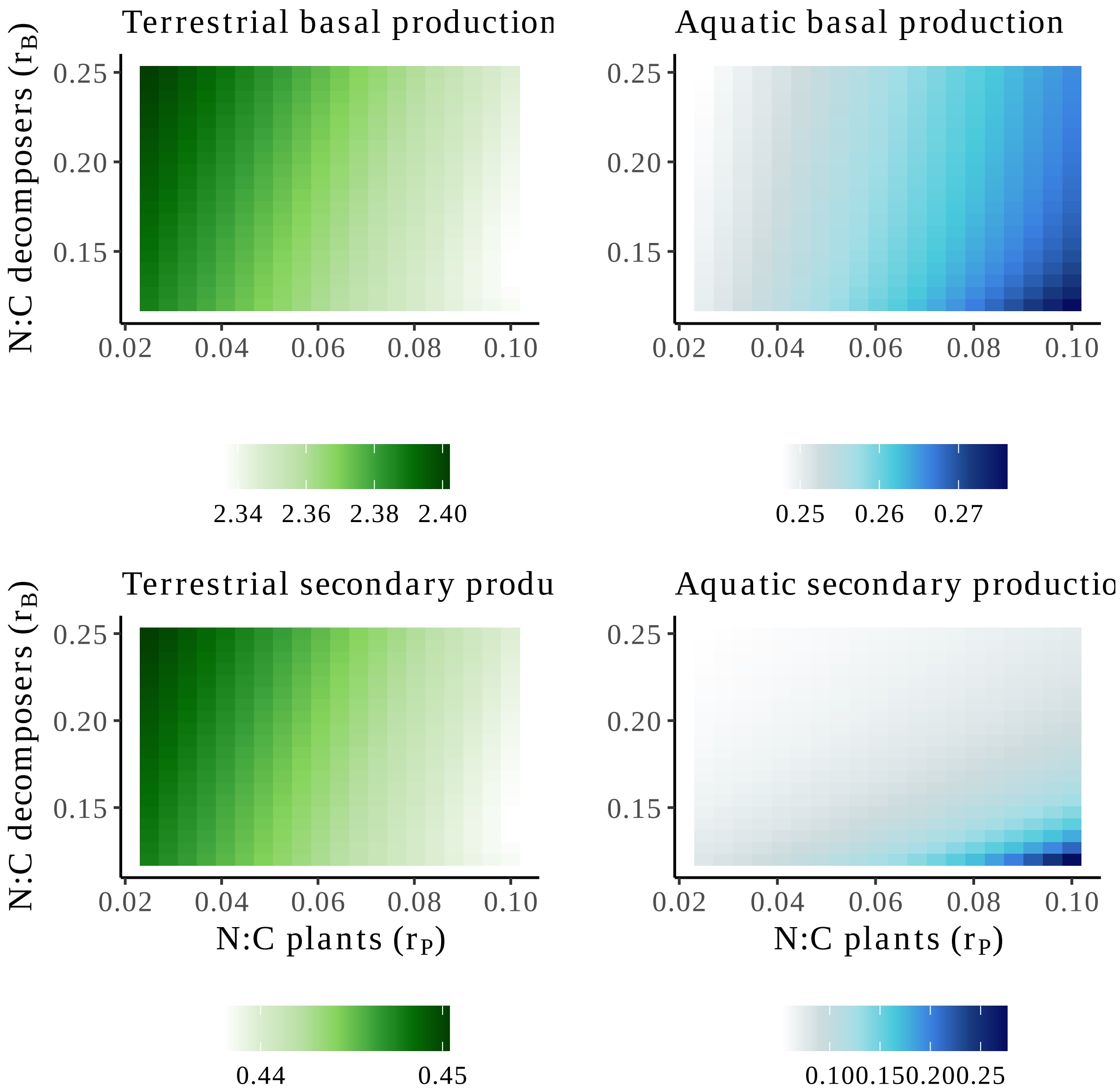


Figure S6.2: **Main conclusions are robust to donor-control functional responses under nitrogen limitation of decomposers.**
The legend is the same as in Fig. S6.1 except that here, decomposers are nitrogen-limited.

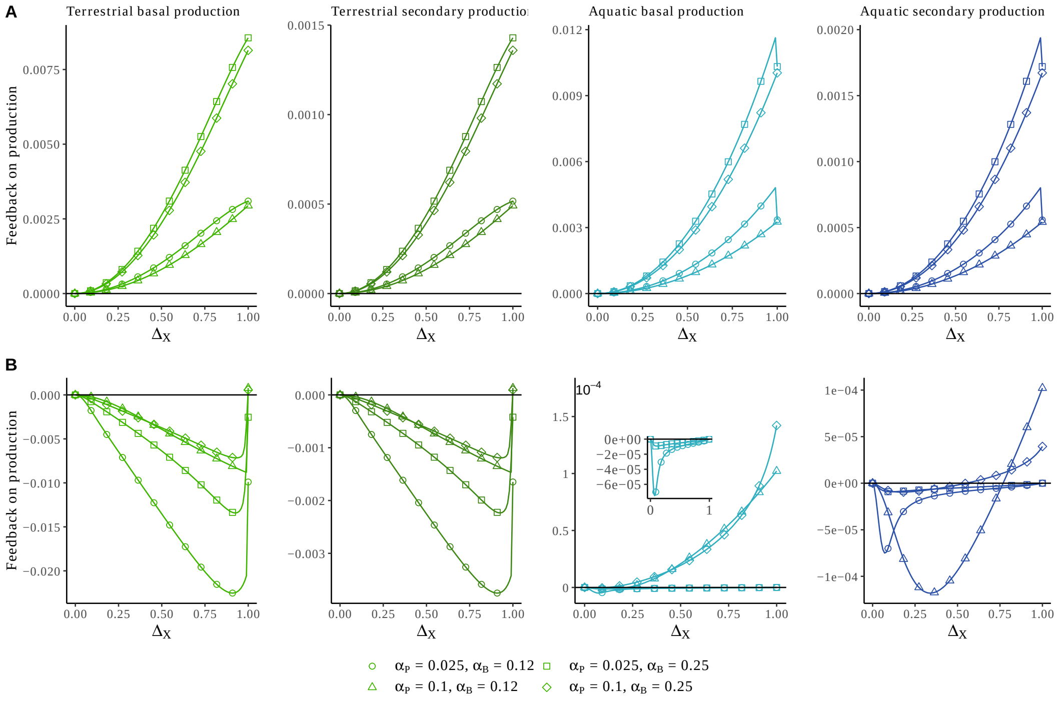


Figure S6.3: **Feedback under both N-limited and C-limited decomposers with donor control functional responses.**

We computed the feedback as shown in Fig. S1.1 and Eq. 4. The feedback is computed for each of the trophic levels in both ecosystems (columns) and under C-limited (A) and N-limited (B) decomposers. The insert shows the sign of the feedback on secondary production for $\alpha_P = 0.1, \alpha_B = 0.25$ (diamond shape) and $\alpha_P = 0.025, \alpha_B = 0.25$ (square shape).

### S7 Sensitivity analysis on the parameter values

In this section, we aim to perform a sensitivity analysis on the parameter value. As the model contains many parameters, we first determined which model parameters were the most sensitive and we further explored for these parameters, whether they qualitatively changed the patterns found in Fig. 3. We performed a sensitivity analysis on the parameter values by varying each parameter independently (see Table. S7.1). For the first step, we varied independently each parameter independently. The range of each parameter under carbon or nitrogen limitation was determined so that *(i)* the simulations remained in the same resource limitation and that *(ii)* the two trophic levels in both ecosystems coexisted (Table S7.1). We measured under the two resource limitations (nitrogen and carbon) the average change in the production of each ecosystem defined as the range of variation of production divided by the range of variation of the parameter value (Fig. S7.1). In both nitrogen- and carbon-limitation, $a_P$, $a_H$, $d_H$ and $e_H$ were the most sensitive parameters (Fig. S7.1B-C). Therefore, for these parameters, we display how the production in both the aquatic and terrestrial ecosystem change with variations of these parameters. Overall, the patterns shown in Fig. 3 are robust to the quantitative variations in parameter values. Finally, we varied the strength of coupling of ecosystems $\Delta$ and observed similar patterns as found in the case $\Delta = 1$ in Fig. 3.

Table S7.1: **Range of parameter variation in the sensitivity analysis.**
Note that for $I_{N_{\mathcal{A}}}$ and $I_{D_{\mathcal{A}}}$ the range under C-limitation (left range) and N-limitation (right range) is different.

| Class | Parameter | Range varied |
|---|---|---|
| **Terrestrial ecosystem** | $I_{N_{\mathcal{T}}}$ | 1-15 |
| | $I_{D_{\mathcal{T}}}$ | 1-15 |
| | $l_{D_{\mathcal{T}}}$ | 0.5-5 |
| | $l_{N_{\mathcal{T}}}$ | 0.5-5 |
| | $m_{\mathcal{T}}$ | 0-1 |
| | $e_G$ | 0.2-1 |
| | $a_{PN}$ | 0.1-0.7 |
| | $a_{GP}$ | 0.1-2 |
| | $d_G$ | 0.05-1 |
| | $d_P$ | 0.01-1 |
| **Aquatic ecosystem** | $I_{N_{\mathcal{A}}}$ | 2-20, 1-2 |
| | $I_{D_{\mathcal{A}}}$ | 2-20, 12-18 |
| | $l_{D_{\mathcal{A}}}$ | 0.5-1 |
| | $l_{N_{\mathcal{A}}}$ | 1-2 |
| | $m_{\mathcal{A}}$ | 0-1 |
| | $e_C$ | 0.3-1 |
| | $e_B$ | 0.5-1 |
| | $a_{BD}$ | 0.83-2 |
| | $a_C$ | 0.3-1 |
| | $d_B$ | 0.01-0.5 |
| | $d_C$ | 0.01-0.15 |

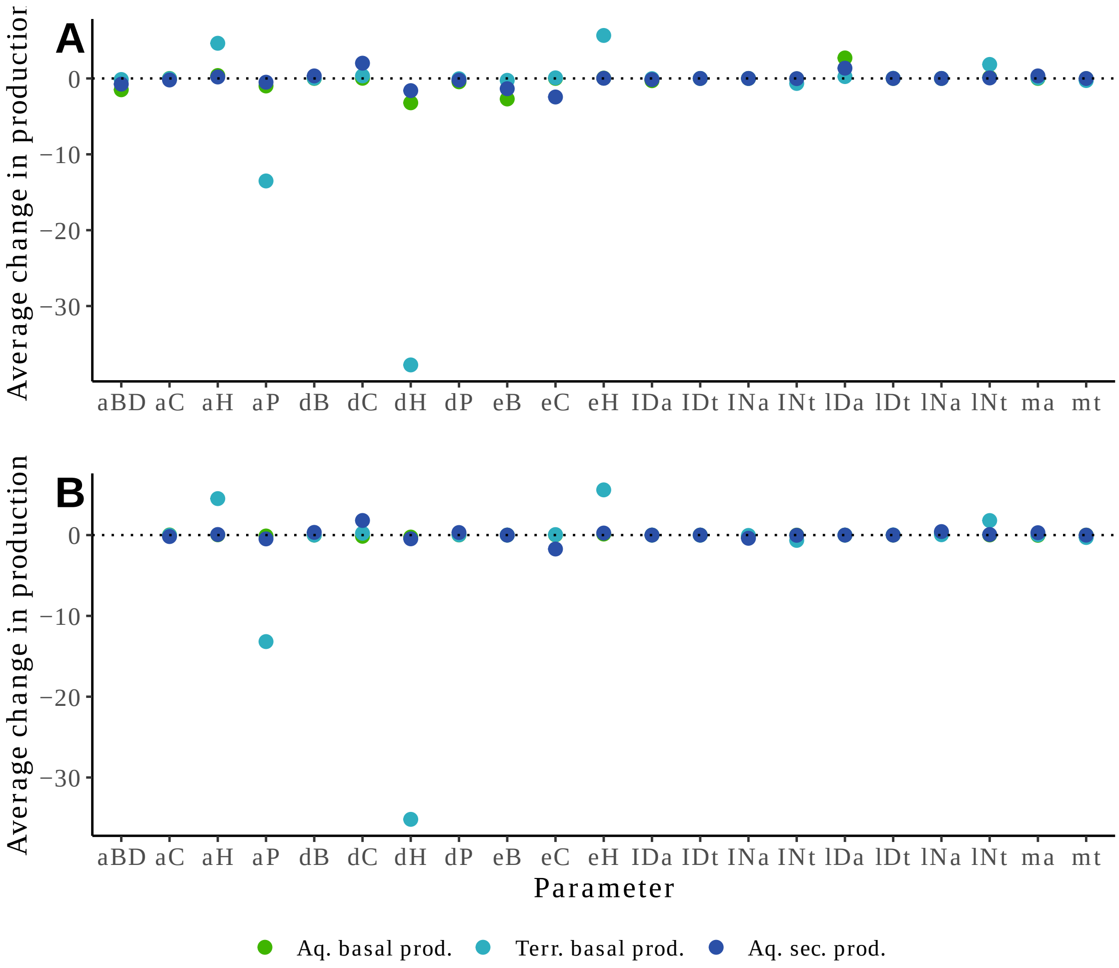


Figure S7.1: **Quantitative sensitivity of ecosystem production to change in parameter values.**
(A & B) We show the average change in the ecosystem production computed as the range of variation of the production divided by the range of variation of each parameter. Each point corresponds to the mean across the 4 values of primary producer stoichiometries. The analyses were performed for (A) C-limited decomposers and (B) N-limited decomposers. Note that we excluded the secondary production in the terrestrial ecosystem as it is qualitatively similar to the primary production. terr. = terrestrial, aq. = aquatic and prod. = production. Some parameters such as $d_H$ under carbon and nitrogen limitation, have a strong quantitative influence on the ecosystem production.

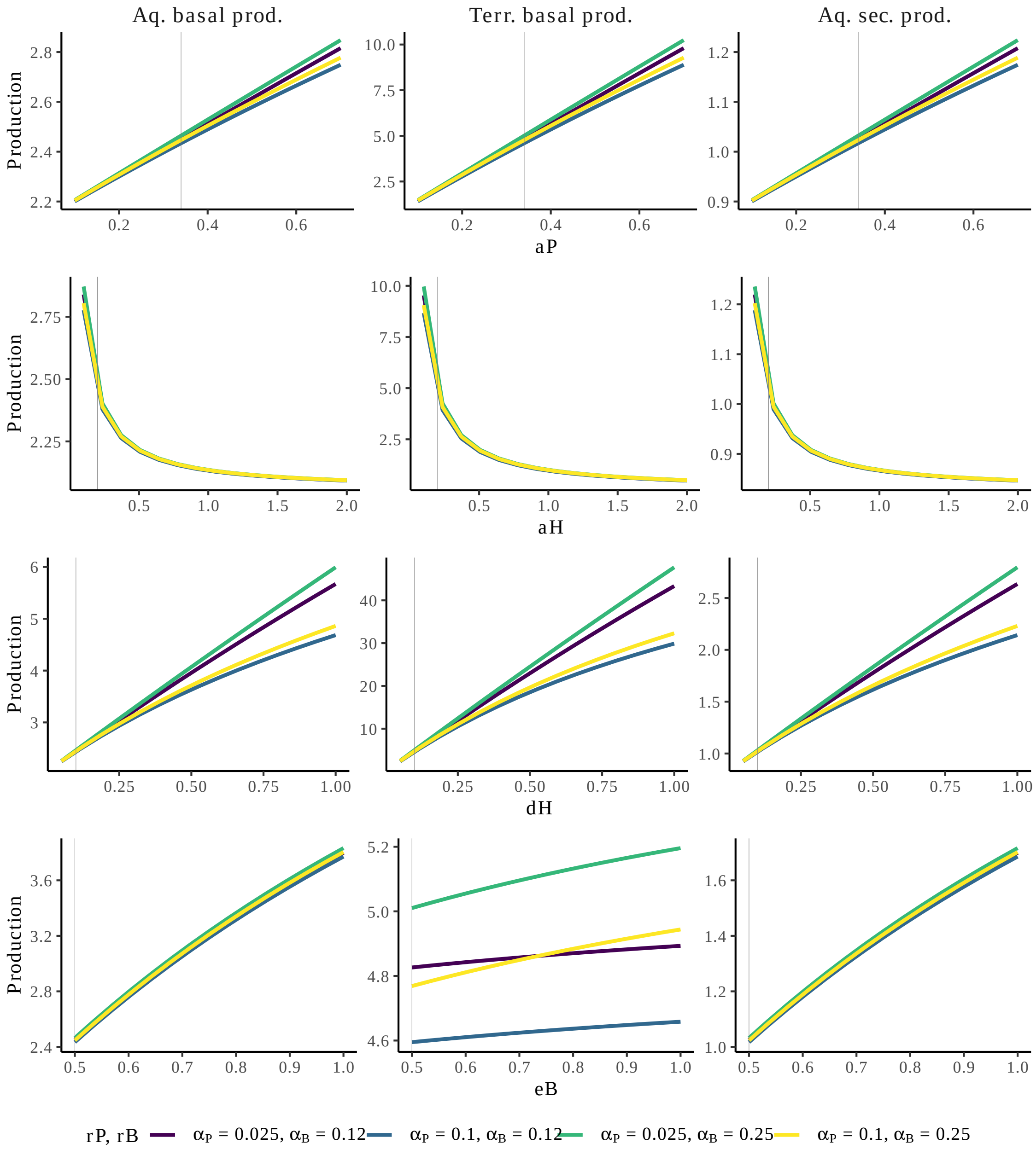


Figure S7.2: **Results are qualitatively robust to parameter variation in C-limitation.** While the most sensitive parameters change quantitatively the production of ecosystems, they do not change the patterns observed in Fig. 3 (*i.e.*, as seen by the relative position of the lines). Here decomposers are carbon-limited.

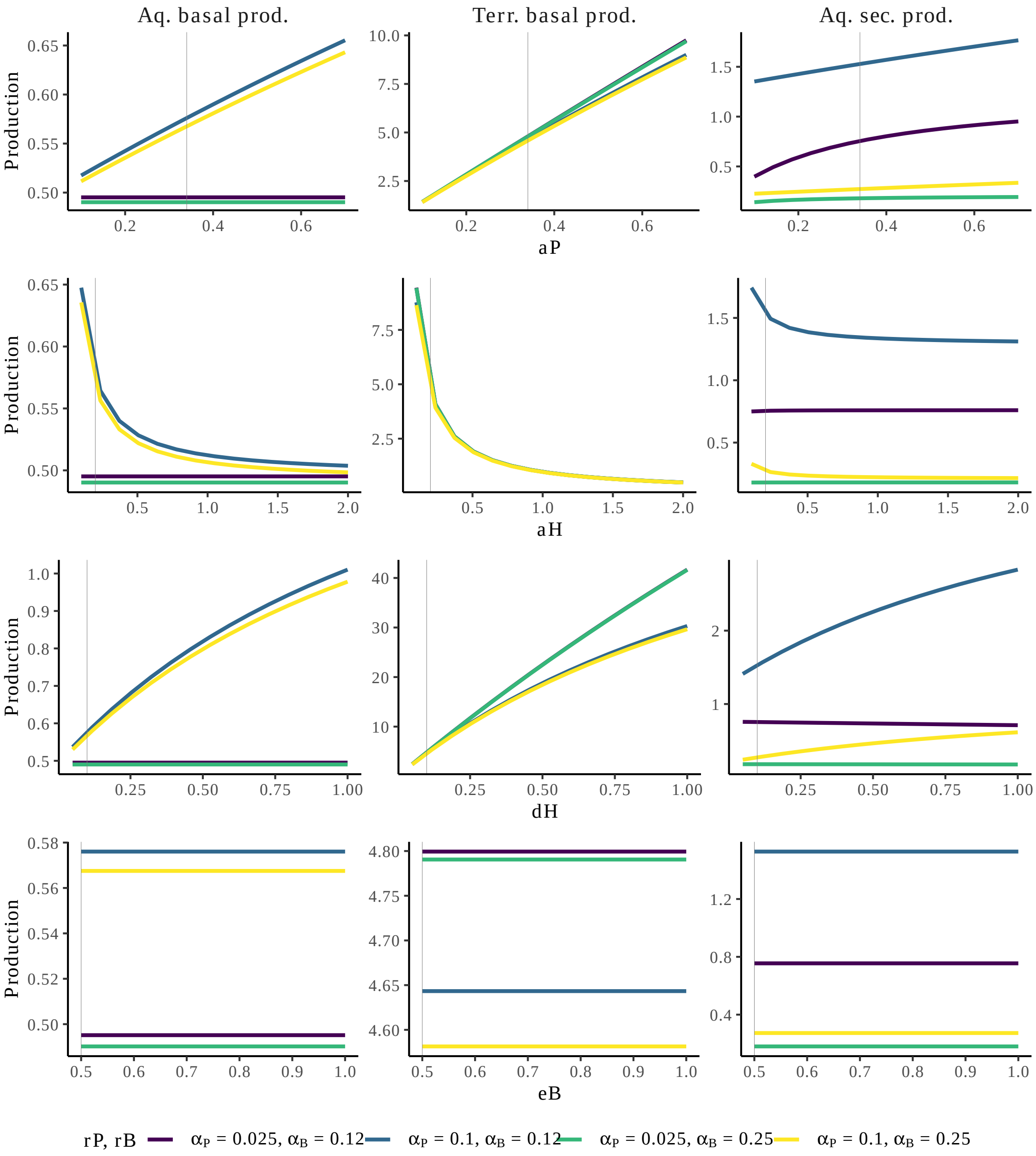


Figure S7.3: **Results are qualitatively robust to parameter variation in N-limitation.** While the most sensitive parameters change quantitatively the production of ecosystems, they do not change the patterns observed in Fig. 3 (*i.e.*, as seen by the relative position of the lines). Here decomposers are nitrogen-limited.

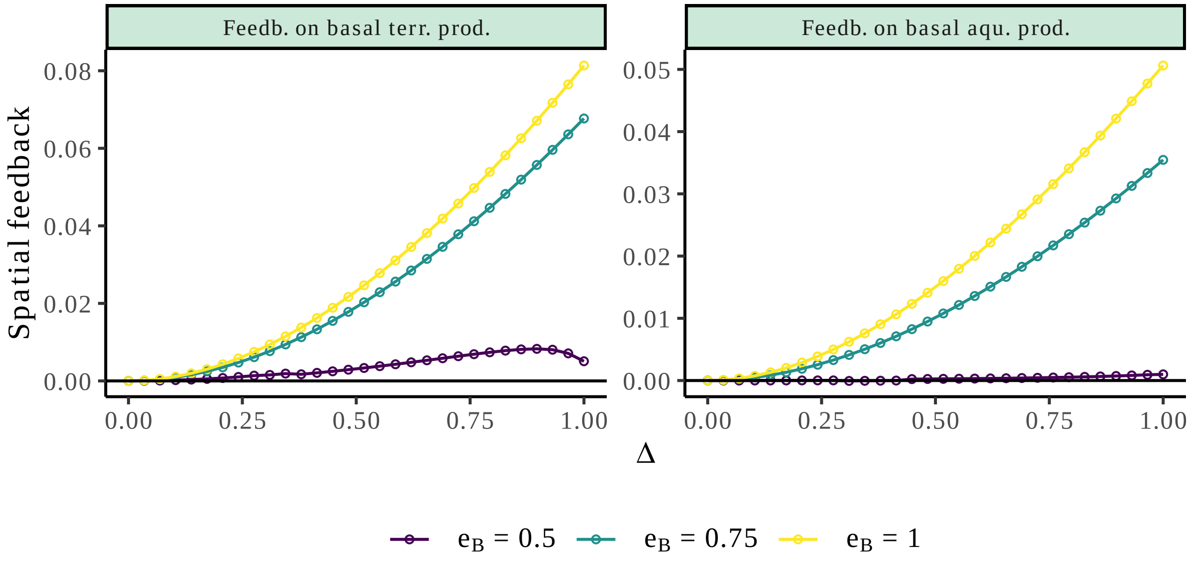


Figure S7.4: **Influence of the growth efficient on the spatial feedback in C-limited aquatic ecosystems.**
We show how the growth efficiency of the decomposers modulates the spatial feedbacks on the basal production of both ecosystems. We only show the results for the basal production as qualitatively similar results are obtained for secondary production. When the growth efficiency of decomposers decreases, the strength of the positive feedbacks on both ecosystems decreases. Other parameters = $\alpha_P = 0.1, \alpha_B = 0.25$.

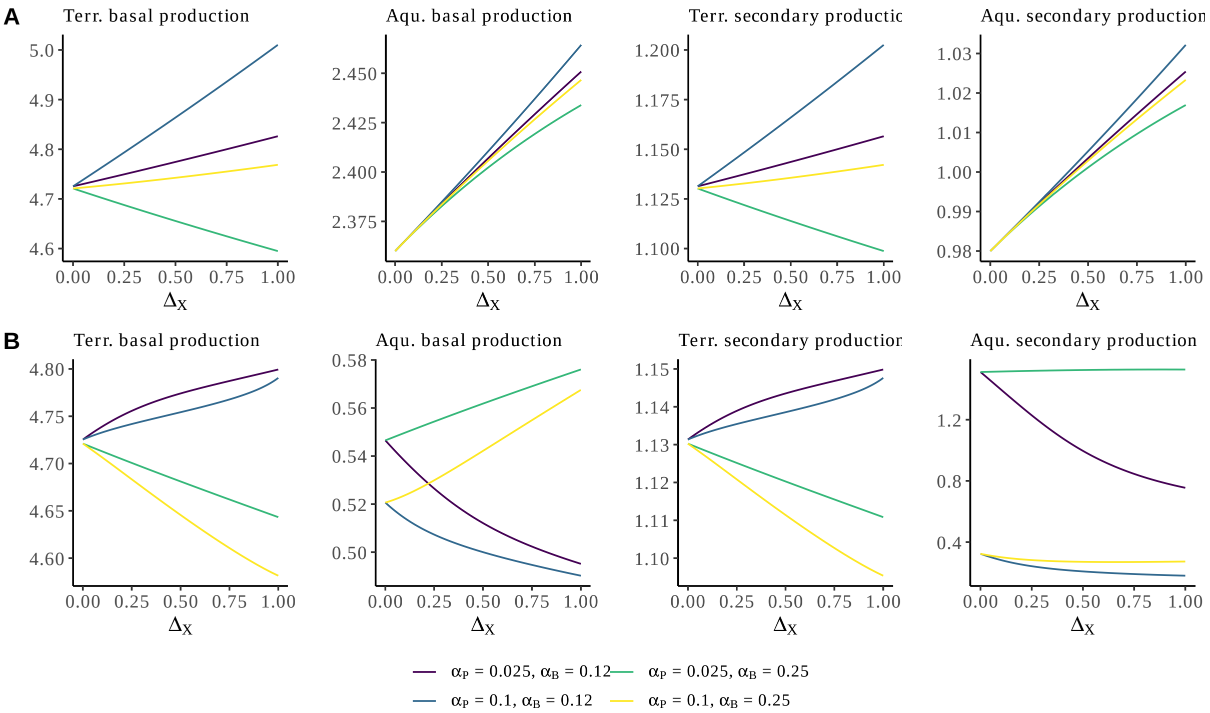


Figure S7.5: **Influence of the strength of ecosystem coupling on the patterns presented in Fig. 3.**
We show the changes in basal and secondary production along the strength of coupling between ecosystems (Δ) for 2 values of plant stoichiometry ($\alpha_P \in [0.025, 0.1]$) and 2 values of decomposers stoichiometry ($\alpha_B \in [0.12, 0.25]$). These cases correspond to the corners of Fig. 3A (top) and Fig. 3B (bottom) along the gradient of coupling between ecosystems. Panel A = C-limited decomposers. Panel B = N-limited decomposers.

### S8 Sensitivity analysis on the asymmetry of flows

We performed a sensitivity analysis to investigate the role of the asymmetry of flows. We focus on the asymmetry of subsidy flow between terrestrial and aquatic ecosystems ($\Delta_{\mathcal{T}} = \{\Delta_P, \Delta_H\}$ *versus* $\Delta_{\mathcal{A}} = \{\Delta_B, \Delta_C\}$). We investigated how differences between $\Delta_{\mathcal{T}}$ and $\Delta_{\mathcal{A}}$ modulate the basal and second production of both ecosystems. The results are displayed in Fig. S8.1. Under the scenario of C-limitation of decomposers, the aquatic ecosystem benefits from higher exports from the terrestrial ecosystem as it relaxes the carbon limitation and fuels the detritus stock (Fig. S8.1A-right). Interestingly for the terrestrial ecosystem, production does not change when detritus from plants and grazers are locally instead of regionally recycled: plants only benefit from nitrogen-rich subsidies exported from the aquatic ecosystem (Fig. S8.1A-left). When decomposers are limited by nitrogen, we see that production in the aquatic ecosystem decreases with increasing subsidies from the terrestrial ecosystem (due to the stoichiometric mismatch mechanism, see main text). Production in the aquatic ecosystem increases when nitrogen-rich detritus from both decomposers and their consumers are locally recycled (Fig. S8.1B-right).

Moreover, we explored how the feedbacks changed depending on the asymmetry of flows by varying $\Delta_{\mathcal{T}}$ (resp. $\Delta_{\mathcal{A}}$) for $\Delta_{\mathcal{A}} \in \{0, 0.25, 0.75\}$ (resp. $\Delta_{\mathcal{T}} \in \{0.25, 0.75\}$) with the stoichiometric parameters taken in Fig. 5. The results are displayed in Fig. S8.2. To simplify the reading, we only display the feedback on the basal production of both ecosystems. Under C-limitation, we observe similar qualitative behavior than when the flows are symmetric (*i.e.*, $\Delta_{\mathcal{T}} = \Delta_{\mathcal{A}}$ in the main text): the feedback is positive and increases with increasing connectivity of ecosystems (Fig. S8.2Aa, Ba colored lines *versus* red line). Interestingly the feedback strength increases when ecosystems are more spatially coupled (higher values of $\Delta_{\mathcal{A}}$ in Fig. S8.2Aa or $\Delta_{\mathcal{T}}$ in Fig. S8.2Ba). This is similar for N-limited decomposers (Fig. S8.2Ab, Bb). To relate to the mechanisms explained in the main text for the N-limitation scenario, we observe that feedbacks are more important when the coupling between ecosystems is more important (as mass-effect increases relatively to stoichiometric mismatch).

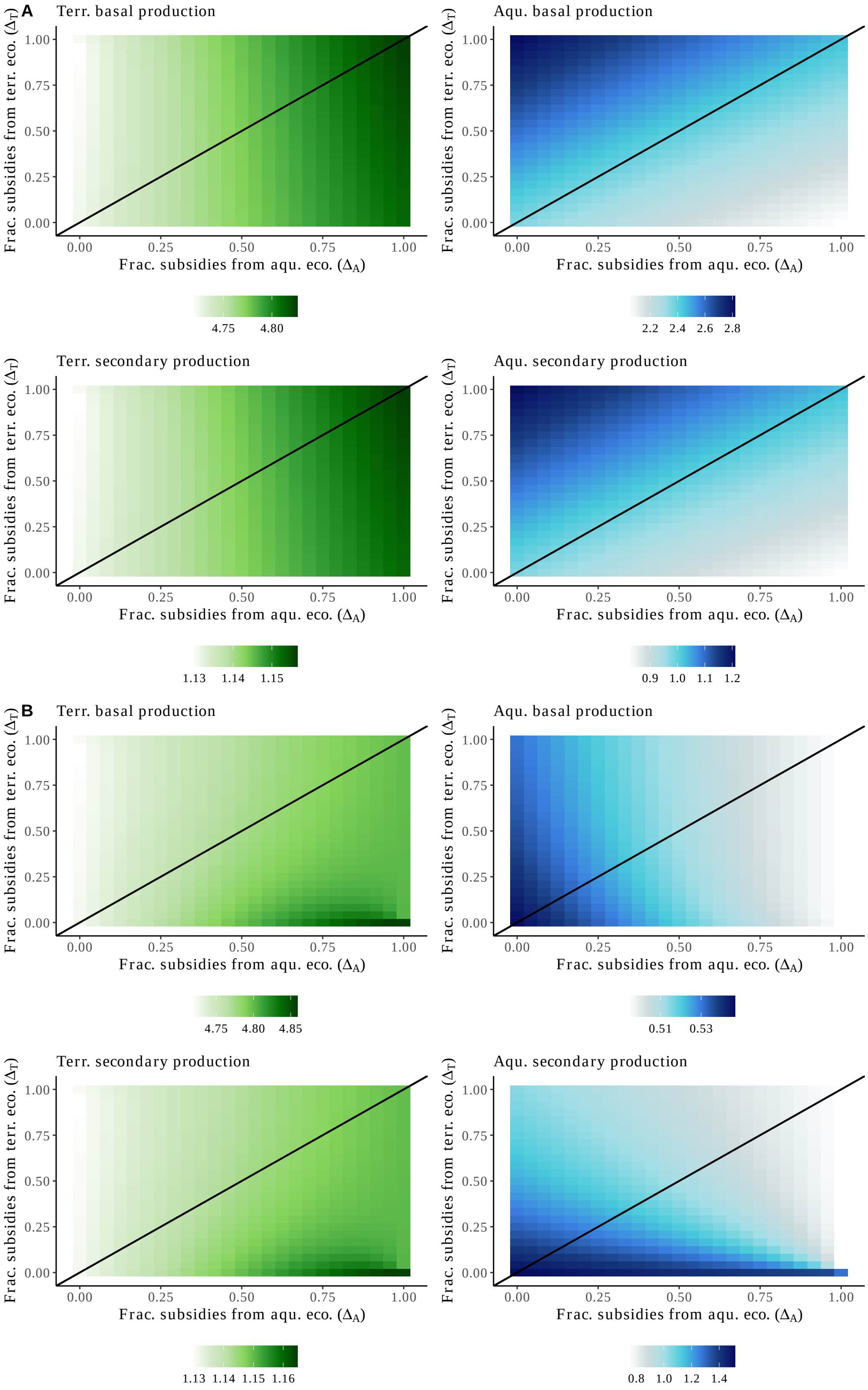


Figure S8.1: **Caption is on next page**

**Fig. S8.1 Sensitivity analysis on the asymmetry of flows on ecosystem production.** We show how basal and secondary productions in both ecosystems are modulated by the asymmetry of subsidies being exchanged at the terrestrial-aquatic ecotone. The black line describes the scenario where the coupling is symmetrical $\Delta_{\mathcal{T}} = \Delta_{\mathcal{A}}$, while below this line (*resp.* above) relatively more subsidies are exported from the aquatic (*resp.* terrestrial) ecosystem. (A) C-limited decomposers. (B) N-limited decomposers. Other parameters: $\alpha_P = 0.025, \alpha_B = 0.12$. Qualitatively similar behaviour is obtained for different combinations of stoichiometries of decomposers and plants. Frac. = Fraction, aqu.= aquatic, terr. = terrestrial, eco. = ecosystem.

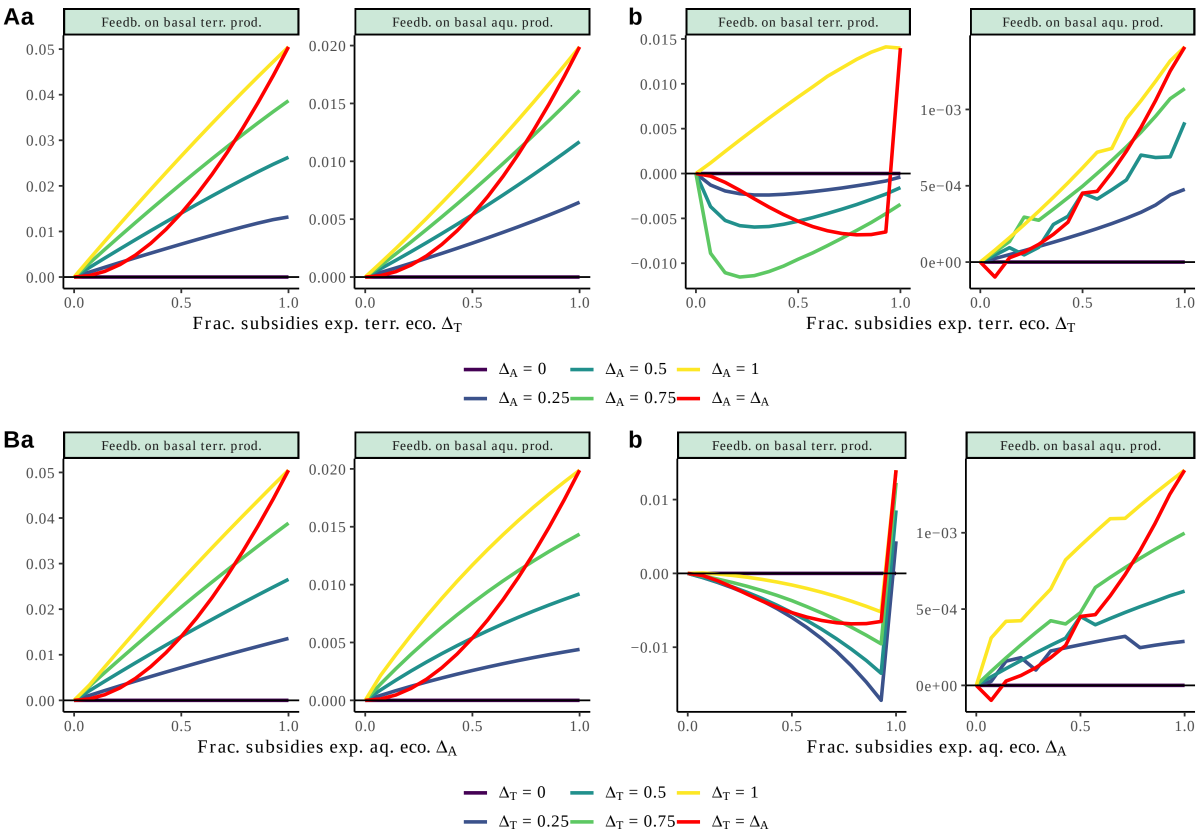


Figure S8.2: **Sensitivity analysis on the asymmetry of flows on spatial feedbacks.** We show how the spatial feedbacks on the basal ecosystem productions are modulated by the asymmetry of subsidies being exchanged at the terrestrial-aquatic ecotone. The black line delimits the negative from the positive feedbacks. (A) We varied the fraction of subsidies exported from the terrestrial ecosystem ($\Delta_{\mathcal{T}}$, x-axis) under different symmetry/asymmetry of flows (colors, red line shows the case of symmetrical coupling as in Fig. 5), and two scenarios of decomposer limitation: C-limited on the left (a) and N-limited on the right (b). Panel B = Similar to (A) but we varied the fraction of subsidies exported from the aquatic ecosystem. Feedb. on basal terr. prod. = Feedback on basal terrestrial production. Feedb. on basal aqu. prod. = Feedback on basal aquatic production.

### Supplementary material references

Attayde, J. L. & Ripa, J. (2008). The coupling between grazing and detritus food chains and the strength of trophic cascades across a gradient of nutrient enrichment. Ecosystems, 11, 980–990.

Boit, A., Martinez, N., Williams, R. & Gaedke, U. (2012). Mechanistic theory and modelling of complex food-web dynamics in Lake Constance. Ecology letters, 15, 594–602.

Buchkowski, R. W., Leroux, S. J. & Schmitz, O. J. (2019). Microbial and animal nutrient limitation change the distribution of nitrogen within coupled green and brown food chains. Ecology, 100, e02674.

Cherif, M. & Loreau, M. (2007). Stoichiometric constraints on resource use, competitive interactions, and elemental cycling in microbial decomposers. The American Naturalist, 169, 709–724.

Cherif, M. & Loreau, M. (2013). Plant–herbivore–decomposer stoichiometric mismatches and nutrient cycling in ecosystems. Proceedings of the Royal Society B: Biological Sciences, 280, 20122453.

Cleveland, C. C. & Liptzin, D. (2007). C:N:P stoichiometry in soil: is there a "Redfield ratio" for the microbial biomass? Biogeochemistry, 85, 235–252.

Danger, M., Daufresne, T., Lucas, F., Pissard, S. & Lacroix, G. (2008). Does Liebig's law of the minimum scale up from species to communities? Oikos, 117, 1741–1751.

Daufresne, T., Lacroix, G., Benhaim, D. & Loreau, M. (2008). Coexistence of algae and bacteria: a test of the carbon hypothesis. Aquatic Microbial Ecology, 53, 323–332.

del Giorgio, P. A. & Cole, J. J. (1998). Bacterial growth efficiency in natural aquatic systems. Annual Review of Ecology and Systematics, 29, 503–541.

Elser, J. J., Fagan, W. F., Denno, R. F., Dobberfuhl, D. R., Folarin, A., Huberty, A., Interlandi, S., Kilham, S. S., McCauley, E., Schulz, K. L., Siemann, E. H. & Sterner, R. W. (2000). Nutritional constraints in terrestrial and freshwater food webs. Nature, 408, 578–580.

Halvorson, H. M., Sperfeld, E. & Evans-White, M. A. (2017). Quantity and quality limit detritivore growth: mechanisms revealed by ecological stoichiometry and co-limitation theory. Ecology, 98, 2995–3002.

Harpole, W. S., Ngai, J. T., Cleland, E. E., Seabloom, E. W., Borer, E. T., Bracken, M. E., Elser, J. J., Gruner, D. S., Hillebrand, H., Shurin, J. B. & Smith, J. E. (2011). Nutrient co-limitation of primary producer communities: Community co-limitation. Ecology Letters, 14, 852–862.

Martinson, H., Schneider, K., Gilbert, J., Hines, J., Hambäck, P. & Fagan, W. (2008). Detritivory: Stoichiometry of a neglected trophic level. Ecological Research, 23, 487–491.

Mcgroddy, M. E., Daufresne, T. & Hedin, L. O. (2004). Scaling of C:N:P stoichiometry in forests worldwide : implications of terrestrial Redfield-type ratios. 85, 12.

Sperfeld, E., Martin-Creuzburg, D. & Wacker, A. (2012). Multiple resource limitation theory applied to herbivorous consumers: Liebig's minimum rule vs. interactive co-limitation: Co-limitation theory applied to herbivores. Ecology Letters, 15, 142–150.

Sperfeld, E., Raubenheimer, D. & Wacker, A. (2016). Bridging factorial and gradient concepts of resource co-limitation: towards a general framework applied to consumers. Ecology letters, 19, 201–215.

Wirtz, K. W. & Kerimoglu, O. (2016). Autotrophic Stoichiometry Emerging from Optimality and Variable Co-limitation. Frontiers in Ecology and Evolution, 4.

Zelnik, Y. R., Manzoni, S. & Bommarco, R. (2021). Primary productivity in subsidized green-brown food webs.

Zou, K., Thébault, E., Lacroix, G. & Barot, S. (2016). Interactions between the green and brown food web determine ecosystem functioning. Functional Ecology, 30, 1454–1465.
